# A computational principle of habit formation

**DOI:** 10.1101/2024.10.12.618033

**Authors:** Kaushik Lakshminarasimhan

## Abstract

Actions are influenced by multiple decision-making systems – including a goal-directed system that favors rewarded actions and a habit system that repeats past actions – but precisely when one system prevails is not known. We show that when competition between these systems is resolved by a winner-take-all mechanism, the precise condition for the emergence of habits can be cast in terms of the well-known probability matching principle. The theory embodies a trade-off in which exploitation, or overmatching, maximizes reward but strengthens habits, while paradoxically, exploration preserves goal-directed behavior by sacrificing rewards. This tradeoff can be averted if learning operates on abstract latent state representations whereby knowing the broader context allows for switching between two habits instead of avoiding one, thus maximizing rewards without forfeiting flexibility. The theory explains a range of animal behaviors as well as task-dependent effects of striatal manipulation, and suggests that neural mechanisms governing exploration implicitly control arbitration between decision-making systems.

## 2 Introduction

We frequently rely on habits, sometimes at the expense of achieving our goals — like when darting out of an elevator and ending up in the wrong hallway. Different neural systems govern goal-directed and habitual decision-making (Yin et al., 2004, 2005; Frank et al., 2009; Balleine & O’Doherty, 2010; Dolan & Dayan, 2013; Wood & Rünger, 2016), with disruptions in the balance between these systems linked to diseases like addiction (Everitt & Robbins, 2005) and obsessive-compulsive disorder (Gillan et al., 2011). However, the exact conditions that cause one system to dominate over the other remain unclear.

A widely held view maps goal-directed and habitual forms of decision-making to prospective and retrospective control systems respectively (Dolan & Dayan, 2013), with behavior governed by the system whose estimate is more reliable (Daw, Niv, & Dayan, 2005). However, in this framework, the two systems are typically modeled using qualitatively different algorithms – model-based and model-free – making their reliability estimates subject to additional assumptions about factors like working memory and computational noise which primarily affect model-based computations. This issue was recently tackled by computational approaches that revived a traditional perspective, modeling learning in both systems using analogous algorithms (Miller, Shenhav, & Ludvig, 2019; Bogacz, 2020). Based on Thorndike’s laws of *effect* and *exercise* (Thorndike, 1911; Guthrie, 1935), learning in the goal-directed and habit systems were proposed to be guided by prediction errors that reinforce *rewarded* actions and *past* actions respectively. In other words, goal-directed and habitual behaviors reflect value-based and value-free, rather than model-based and model-free, computations, and aligns with the motivational perspective of control in which goal-directed and habitual actions are driven by intrinsic values and extrinsic cues respectively (Ceceli & Tricomi, 2018). Studies employing models of this class have provided a compelling account of the emergence of habits from extended training (Miller et al., 2019; Bogacz, 2020; Greenstreet et al., 2022) but have not yet uncovered the criteria for habit formation.

Experiments show that training duration does not necessarily predict whether a behavior is habitized (Robbins & Costa, 2017; Garr et al., 2021; LaFlamme et al., 2022). Clearly, no amount of training would produce habits if actions are chosen randomly by ignoring rewards. One possibility is that whether habits form after extended training depends on the relation between the statistics of rewards and actions during learning. This relationship has been examined extensively through the lens of probability matching (Gallistel, 1993). In discrete-trial tasks such as the bandit problem, behavior is said to obey probability matching when the likelihood of choosing an action matches the likelihood of being rewarded for that action. This is in fact a suboptimal strategy because rewards are maximized only if the action associated with a higher reward probability is chosen exclusively (Mongillo, Shteingart, & Loewenstein, 2014). Yet, animals exhibit probability matching in many cases, revealing a tendency to systematically underexploit (Grant, Hake, & Hornseth, 1951; Bullock & Bitterman, 1962; Bitterman, 1971; Vulkan, 2000; Rubinstein, 2002; Koehler & James, 2014; Kubanek & Snyder, 2015). This puzzling finding is usually taken to reflect wrong internal models (Yu & Huang, 2014), different timescales of learning (Iigaya et al., 2019) or resource-constrained computations (Vul, Goodman, Griffiths, & Tenenbaum, 2014; Kubanek & Snyder, 2015; Bari & Gershman, 2023), overlooking their impact on habits. In this study, we analyze the circumstances under which habits form by examining the implications of the exploration-exploitation balance on learning.

To do so, we developed and analyzed an analytically tractable model of learning and decision-making in which the competition between goal-directed and habit systems is categorically resolved through a winner-take-all mechanism instead of soft arbitration. This revealed a computational principle linking probability matching and habits whereby overmatching (or exploitation) promotes the formation of rigid habits while exploration sustains flexible, goal-directed behavior. We show how this exploration-exploitation trade-off could be circumvented by learning on abstract latent state representations. The theory accounts for a range of experimental findings and offers a quantitative framework to study the neural basis of goal-directed and habitual decisions by factoring inter-individual variability in exploratory behavior.

## 3 Results

Consider an environment in which choosing an action *a* in state *s* yields some reward *r*. To analyze when habits are formed, we construct a model comprising two parallel learning systems – a goal-directed system and a habitization system – that estimate different statistics (**Figure 1A**). For each state *s*, the goal-directed system uses past rewards to estimate the expected reward associated with action *a* i.e., *action value*, denoted by *Q_s,a_*, while the habitization system uses past actions to estimate the tendency of taking action *a* i.e., *action preference*, denoted by *H_s,a_*. Following previous work, we assume that learning in these systems is mediated by reward prediction error, *δ_r_* = *r* − *Q_s,a_*, and action prediction error, *δ_a_* = 1 − *H_s,a_*, respectively such that Δ*Q_s,a_* ∝ *δ_r_* and Δ*H_s,a_* ∝ *δ_a_* when choosing action *a* in state *s* (Miller et al., 2019; Bogacz, 2020; Greenstreet et al., 2022). Preferences for both the chosen and non-chosen action are updated to ensure unbiased learning in the habitization system (Methods – Equation 4).

**Figure 1:**
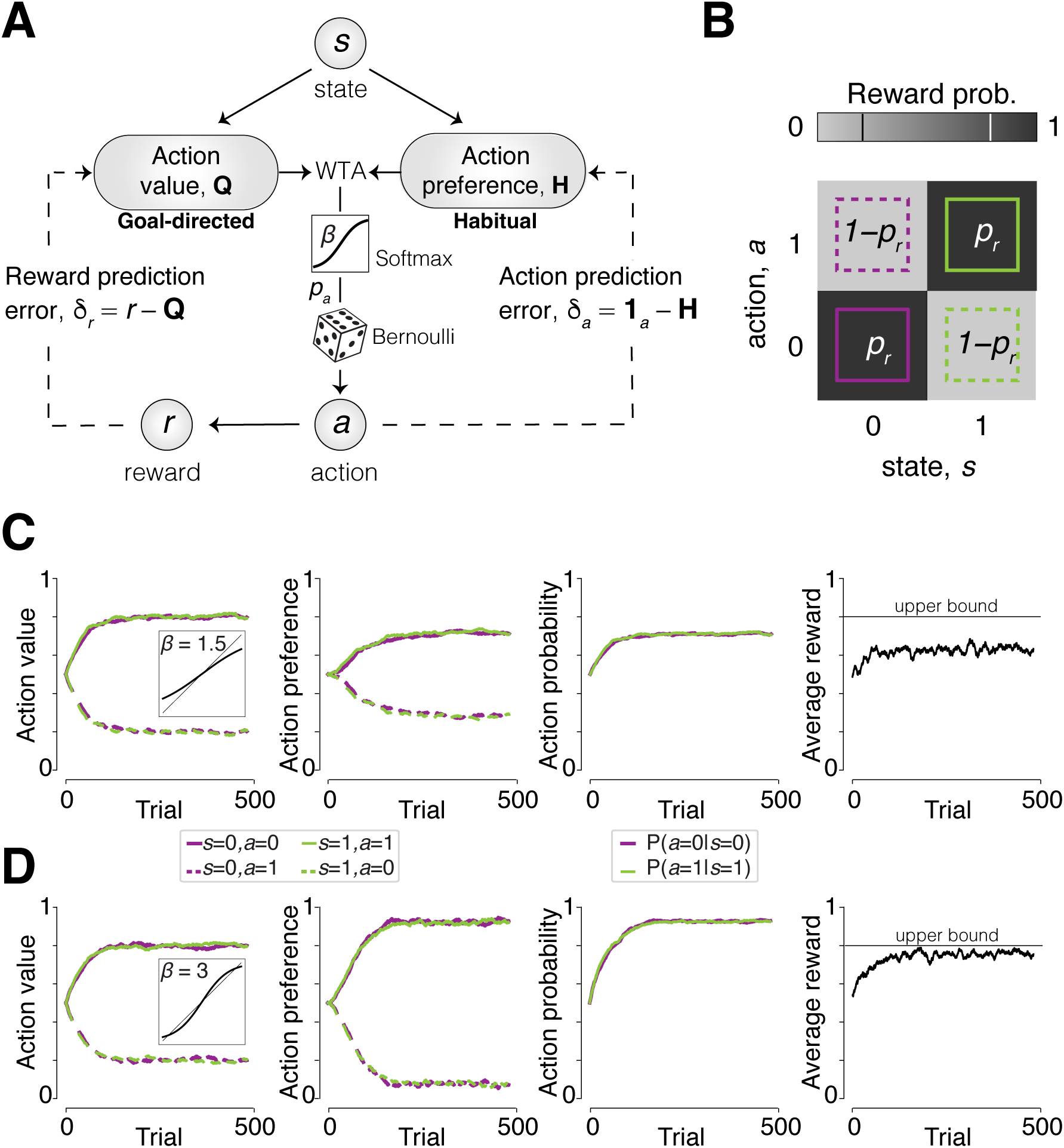
Model schematic and simulation. **A.** Illustration of the model with dual systems. Solid lines indicate the transformations during the decision-making process and dashed lines indicate error signals that mediate the learning process. **B.** Reward probabilities for different states and actions in a binary decision-making task. **C.** Evolution of the goal-directed system’s action value estimate (*Q*), the habitization system’s action preference estimate (*H*), action probability (*p_a_*), and average reward (*r*) when learning a task in which *p_r_* = 0.8 and the degree of exploitation is low (*β* = 1.5). The horizontal line in the rightmost panel denotes the maximum possible average reward for an optimal policy (*r*_max_ = *p_r_* = 0.8). **D.** Similar to the panels in **C** but when the degree of exploitation is relatively high (*β* = 3).

Prior works assumed that the decision variable is computed by taking a weighted sum of the estimates from the two systems. Because our objective here is to determine the conditions under which one system prevails over the other, we instead assume a winner-take-all (WTA) mechanism that chooses the system that is more greedy in any given state *s* by comparing the vectors *Q_s_* and *H_s_* (Methods – Equation 5). Following standard approaches, actions are then selected based on the probability distribution obtained by applying a policy mapping function e.g., the softmax operator on the estimates of the chosen system. In case of binary actions, this means *a* ∼ Bernoulli(*p_a_*) where the probability of choosing action *a* is given by:

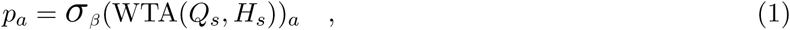

where *σ_β_* (·) denotes the softmax operator parameterized by *β* which controls the exploration-exploitation trade-off (Methods – Equation 6). Smaller values of *β* result in more exploration while larger values lead to exploitation. We will now show that this degree of freedom plays a crucial role in determining whether actions eventually become habitual or goal-directed after learning.

### 3.1 Relation between habits and probability matching

We begin by simulating the model on a task in which states, actions, and rewards are all binary. Rewards are stochastic such that in each state, one action is rewarded with probability *p_r_* while the other action has a reward probability of 1 − *p_r_* (**Figure 1B**). In this setting, both value *Q* and preference *H* can themselves be interpreted as probabilities (of receiving a reward and choosing an action respectively), enabling us to directly compare their magnitudes to determine the winner (Supplemental Figure S1A). We found that when *β* is small, the model is driven by the goal-directed system in steady state i.e., when learning has plateaued, and the harvested reward is substantially smaller than the upper bound *p_r_* (**Figure 1C**; Supplemental Figure S1B). In contrast, when *β* is large, the model is eventually driven by the habitization system and the reward is nearly optimal (**Figure 1D**). This simulation hints at the possibility that habits are formed when the degree of exploitation exceeds a critical value.

Indeed, a simple calculation reveals a precise condition for habit formation. As learning progresses, the estimate associated with the most rewarding action approaches the reward probability asymptotically in the goal-directed system (*Q_s,a_* = *p_r_*) (Methods – Equation 7) while the corresponding estimate in the habitization system approaches the action probability (*H_s,a_* = *p_a_*) (Methods – Equation 8), yielding:

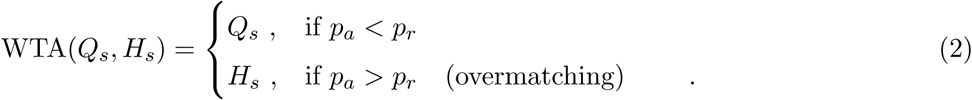

Thus, decisions are goal-directed as long as the probability of choosing the best action does not exceed the probability of reward associated with it. On the other hand, habits are formed when overmatching. Note that although action probabilities depend on the specific operator that transforms the decision variable into actions, the above principle holds universally for all operators. In case of the softmax operator, we can compute the critical level of exploitation *β* = *β*_crit_ that corresponds to probability matching behavior (Methods – Equation 9). Action probabilities in the two regimes can be estimated by combining equations 1 and 2 which gives:

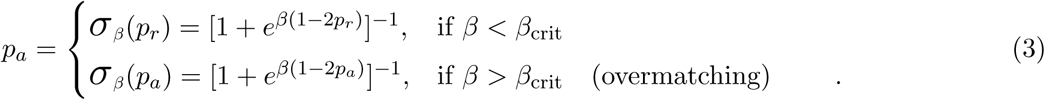

This result encapsulates the fundamental dichotomy between goal-directed and habitual actions, which can be understood by examining how actions differ when *β* is smaller or larger than *β*_crit_ for the example simulated in **Figure 1C**,**D**. When *β* is smaller, action probability is less than the reward probability and critically, depends on the latter which is characteristic of goal-directed behavior (**Figure 2A** – left). When *β* is larger, action probability is greater than the reward probability (overmatching) and, as typical of habits, is independent of the latter (**Figure 2A** – right). Instead, the frequency of actions in this regime is determined by the fixed point equation *p_a_* = *σ_β_* (*p_a_*) and thus depends only on the level of exploitation. This is more explicit when directly looking at action probability as a function of reward probability, which shows that actions vary smoothly with rewards in the goal-directed regime but are stereotypical in the habitual regime (**Figure 2B**).

**Figure 2:**
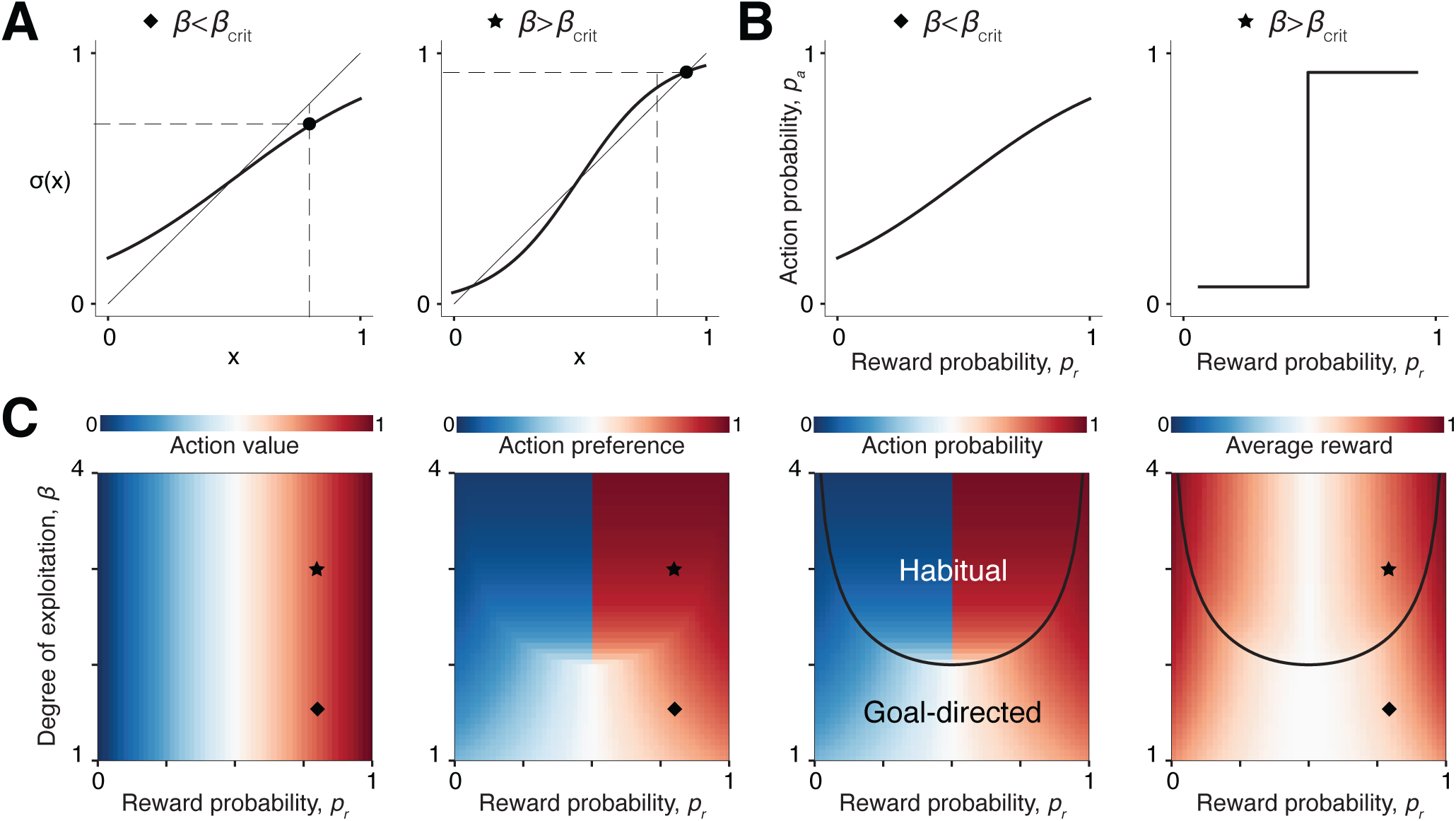
Overmatching promotes habit formation. **A.** Softmax function, *σ*(·) with parameter *β* = 1.5 (left) and *β* = 3 (right). Vertical dashed lines denote the reward probability, *p_r_*=0.8. Horizontal dashed lines denote the probability of choosing the rewarding action, *p_a_* after learning. **B.** Probability of choosing an action as a function of reward probability in steady state i.e., after learning has converged. **C.** Action value (*Q*, left), action preference (*H*, left middle), action probability (*p_a_*, right middle), and average reward *r* (right) as a function of reward probability *p_r_* and the degree of exploitation *β* in steady state. Symbols (star and diamond) correspond to the parameter settings in panel **A**. Black curve corresponds to the critical level of exploitation *β*_crit_ corresponding to matching behavior (*p_a_* = *p_r_*).

We can apply this principle to identify the regime of habits across different levels of reward uncertainty and exploitation. Action value *Q* estimated by the goal-directed system depends only on the level of reward uncertainty (**Figure 2C** – left). In contrast, action preference *H* estimated by the habitization system depends only on the level of exploitation provided it is large enough to form habits (**Figure 2C** – left middle). The critical value of *β* separating goal-directed and habit regimes increases as *p_r_* approaches 0 or 1 (**Figure 2C** – right middle, black curve) implying that the level of exploitation needed to form habits increases as one of the actions becomes overwhelmingly more rewarding. Conversely, habits form more readily when the payoffs of different actions are uncertain. Notably, for any given distribution of rewards, habits are always more rewarding than goal-directed actions (**Figure 2C** – right). This is because the optimal strategy in this setting is to always choose the more rewarding action as stated earlier. However, as we show in a later section, habits could be detrimental when reward statistics change unless learning operates on state representations derived from hierarchical inference.

We emphasize that the above findings are tied to the principle of matching and not to any specific decision rule. Probability matching law can be used to delineate goal-directed and habit regimes even when actions are chosen based on an *ɛ*-greedy rule instead of softmax (Supplemental Figure S2A) or when the arbitration between the two systems is implemented by a weighted sum instead of WTA (Supplemental Figure S2B).

### 3.2 Habits in operant conditioning

The principle can be readily generalized to actions with non-binary and/or asymmetric rewards by expressing the criterion for habit formation in terms of value matching rather than probability matching. Specifically, choosing actions with a ratio greater than their values (reward probability × reward magnitude) should promote habits (Methods – Equation 10; Supplemental Figure S3). This formulation applies to free operant conditioning paradigms historically employed to study habits. In such paradigms, animals get reward *r* for performing an action e.g., lever press and no reward otherwise. While this departs from the binary choice discrete-trial design considered so far, we can apply the theory to this setting by dividing time into discrete bins and assuming the animal gets a small intrinsic unit reward for each bin spent not pressing the lever, representing the pleasure of relaxation. Since the reward schedule is not exactly known to the animal, reward follows any given action with probability *p*. This leads to the criterion that habits form when the proportion of time bins with lever presses exceeds 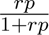 (Methods – Equation 11). As we demonstrate below, whenever uncertainty is high (*p* is low), the above threshold decreases, facilitating habit formation in different operant conditioning paradigms.

First, when obtaining reinforcement requires two sequential actions (e.g., lever press and chain pull, **Figure 3A** – left), the distal action is more vulnerable to habitization (Balleine, Garner, Gonzalez, & Dickinson, 1995). Let *a^D^* and *a^P^* denote the distal and proximal action respectively, and let 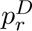 and 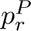 be the corresponding reward probabilities i.e., the proportion of times those actions culminate in reward. The reward probability for the distal action is necessarily smaller than that for the proximal action since, 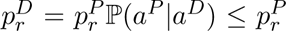 where the inequality arises from failures to complete the sequence. Recall that for any given *β*, the balance shifts in favor of the habit system as reward probability decreases. Therefore if the failure rate is high, the action value, *Q*, of the distal action drops below its action preference, *H*, causing the distal action to become habitized (**Figure 3A** – left middle). While the critical level of exploitation *β*_crit_ needed for habitization decreases with failure rate (Methods – Equation 12), even low failure rates can induce habits if the level of exploitation is moderately high (**Figure 3A** – right middle). Balleine et al. showed that following a reward devaluation procedure, the rate of proximal, but not distal, action decreases. For simplicity, we model the effect of outcome devaluation in this paradigm by scaling down the *Q* value before testing as in previous work (Miller et al., 2019) (Methods). Simulations with failure rate matched to data recapitulated the empirical results (**Figure 3A** – right).

**Figure 3:**
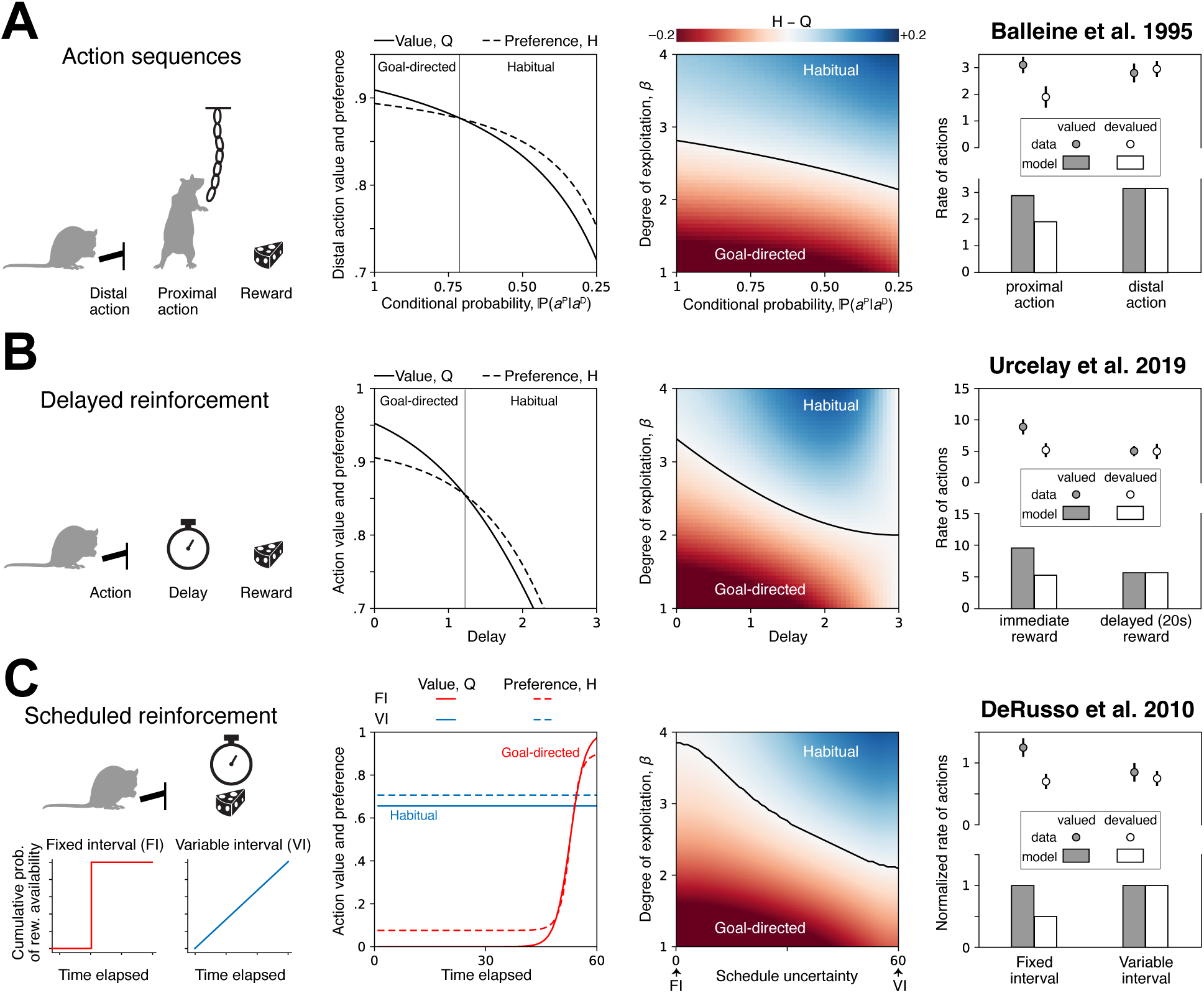
Theory explains habits in operant conditioning paradigms. **A.** Left: Rats perform a lever press and a chain pull in sequence to obtain reward. Left middle: Value and preference of the distal action after learning with *β* = 2.5, as a function of the probability of sequence completion. Right middle: The difference between distal action preference and value after learning, as a function of the probability of sequence completion and the degree of exploitation. Black curve corresponds to the critical level of exploitation *β*_crit_ needed for behavior to become habitual. Right: Number of proximal and distal actions per minute in rat data and model, before and after devaluation. **B.** Left: Rats perform a lever press to obtain a delayed reward. Left middle: Value and preference of the action after learning with *β* = 2.5, as a function of the delay expressed in units of the eligibility trace *τ* (Methods). Right middle: The difference between action preference and value after learning, as a function of the delay and the degree of exploitation. Right: Number of actions per minute for immediate and delayed rewards in rat data and model, before and after devaluation. **C.** Left: Rats perform a sequence of lever presses to obtain a reward delivered according to a FI or VI schedule. Left middle: Value and preference of the action after learning with *β* = 2.5, as a function of the time elapsed since the previous reward (Methods). Right middle: The difference between time-averaged action preference and value after learning, as a function of the schedule uncertainty (Methods) and the degree of exploitation. Right: Normalized number of actions per minute for FI and VI schedules in rat data and model, before and after devaluation.

Second, (Urcelay & Jonkman, 2019) showed that habits are more likely to form when rewards are delayed. Recall that, after learning, estimates in the goal-directed and habit systems converge to expected reward and action, respectively. This requires the timing of prediction error signals that modulate plasticity to coincide with the activation of neurons that encode states and actions (Fisher et al., 2017). This biological constraint cannot be met by reward prediction errors when rewards are delayed. We model this by introducing an eligibility trace that decays exponentially with time constant *τ* taken to be in the order of seconds (**Figure 3B** – left). Delayed rewards bias the estimates of the goal-directed system such that action value is underestimated more as delay gets longer, allowing the habit system to eventually dominate (**Figure 3B** – left middle). The minimum delay *T* for habit formation depends on whether exploitation exceeds the critical value *β*_crit_ (Methods – Equation 13). When *T* ≫ *τ*, low levels of exploitation is sufficient to form habits (**Figure 3B** – right middle). Simulations with *τ* = 10*s* provided a good match to data from Urcelay et al. showing a delay-dependent effect of reward devaluation (**Figure 3B** – right).

Third, we consider a widely reproduced result on habit formation, namely the effect of reinforcement schedule. Behaviors tend to become habitized when reinforcements are delivered on a variable interval (VI) schedule but not on a fixed interval (FI) schedule (DeRusso et al., 2010). Under FI schedule, the reward is baited periodically whereas under VI, the interval between baits is variable (**Figure 3C** – left). Therefore, lever presses towards the end of the interval are strongly reinforced in FI, increasing their value, while earlier actions are strongly devalued (**Figure 3C** – left middle, solid blue). In contrast under VI, learned values are weaker and more uniformly distributed in time (**Figure 3C** – left middle, solid red). Consequently, even for a moderate degree of exploitation, *β*, action preferences dominate over action values in VI schedule (**Figure 3C** – left middle, dashed red). As the schedule shifts from FI to VI on a continuum, the critical degree of exploitation needed to form habits decreases dramatically (Methods; **Figure 3C** – right middle). This explains why habits are readily induced under VI but not in FI schedules. We confirmed this by simulating the effect of reward devaluation which yielded results that qualitatively match the data (**Figure 3C** – right).

### 3.3 Habits in reversal learning

While habits maximize rewards in stationary settings, they can be detrimental in environments where reward contingencies change over time. In fact, as mentioned above, most diagnostic tests for habits are predicated on the idea that habitual actions are insensitive to reward devaluation (Adams, 1982; Dickinson, Balleine, Watt, Gonzalez, & Boakes, 1995; Tricomi, Balleine, & O’Doherty, 2009; Rhodes & Murray, 2013). Another classic paradigm to probe whether animals flexibly adapt to changes in reward statistics is probabilistic reversal learning, where the action leading to a (probabilistic) reward changes abruptly (Methods). As expected, simulating the model to perform this task shows that habits impede behavioral adaptation. Specifically following a reversal, negative reward prediction errors trigger relearning in the goal-directed system. When decisions are governed by this system, the behavioral policy gradually changes in accordance with the revised estimates of action value *Q* and the average reward recovers (**Figure 4A**). In contrast, when decisions are under habitual control, changes in the goal-directed system do not propagate to the decision since the behavioral policy in this case depends on rigid action preference *H*, resulting in persistent errors (**Figure 4B**). Therefore, in its simplest form, the theory predicts that animals that overmatch during the initial acquisition phase will have formed habits and thus cannot adapt to reversals (Supplementary Figure S4A), implying that preserving flexibility in dynamic environments entails forsaking rewards by resisting habit formation.

**Figure 4:**
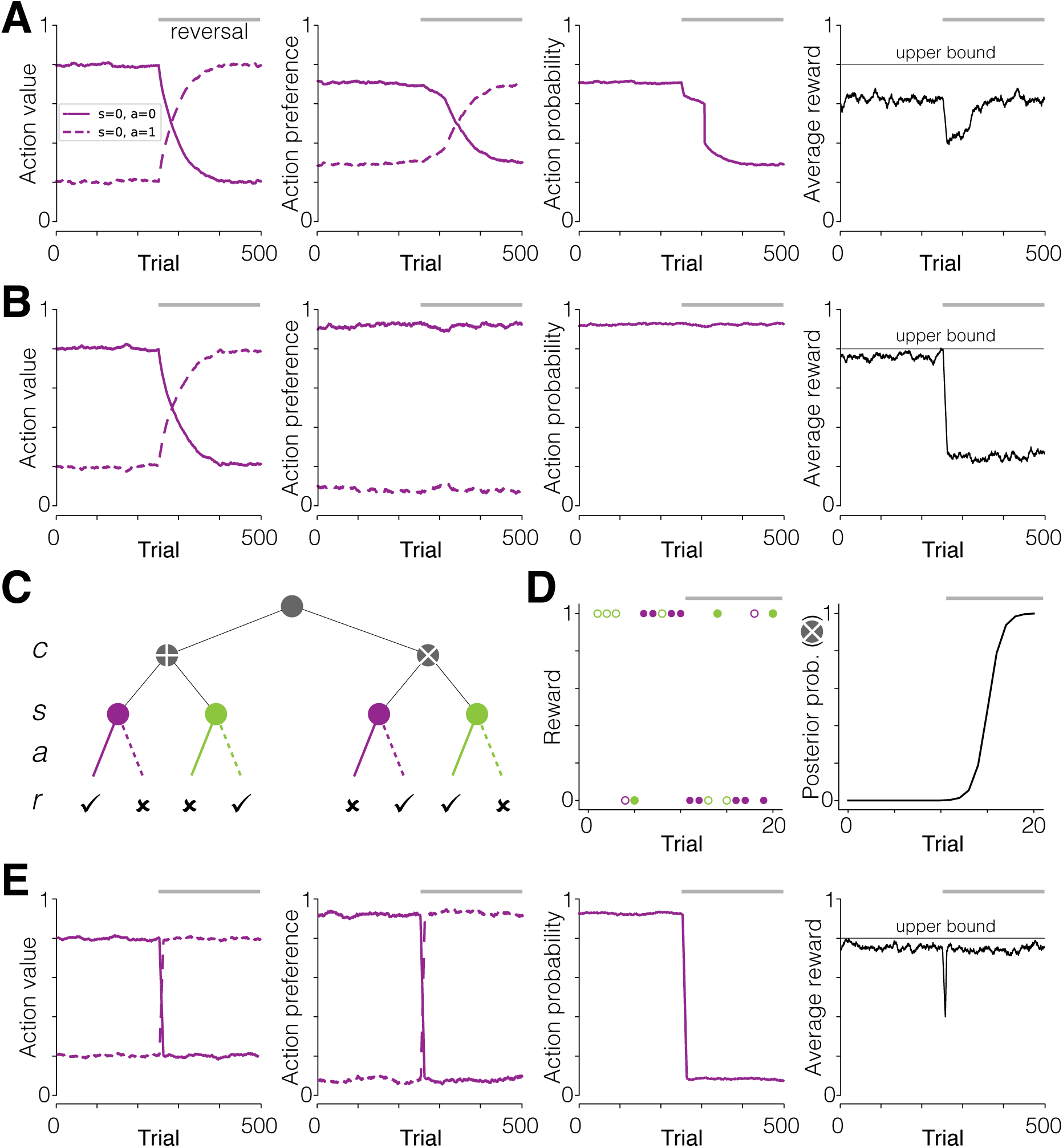
Context inference complements habit formation in probabilistic reversal learning. **A.** Evolution of the action value estimate (*Q*), the action preference estimate (*H*), action probability (*p_a_*), and average reward (*r*) when learning a task in which *p_r_* = 0.8 and the degree of exploitation is low (*β* = 1.5). The horizontal line in the rightmost panel denotes the maximum possible average reward for an optimal policy (*r*_max_ = *p_r_* = 0.8). Note that quantities are plotted only for one of the two states. **B.** Similar to **A**, but when the degree of exploitation is high (*β* = 3). **C.** Structure of the repeated reversal learning task, showing the hierarchical relationship between sensory state *s* and the context *c*. **D.** Example simulation showing actions and rewards around the time of reversal (left) and the associated Bayesian updates in belief about the context (right). **E.** Similar to **B**, but informed by contextual inference.

But a hierarchical view suggests that habitization does not preclude adaptation to reversals. Repeated exposure to both reward contingencies can enable one to identify them as different contexts, and learn separate policies for each context. In other words, the same state, *s* can trigger different habitual actions under different contexts, *c* when learning operates on the augmented representation (*c*, *s*). Here we use state to refer to any low-level latent variable that can be inferred from the stimulus on a given trial, while context is a more abstract latent variable at a higher level of the hierarchy encapsulating reward contingency (**Figure 4C**). Based on the known statistical relationship between states, actions and rewards in the two contexts acquired during training, it is possible to rapidly infer which context is more consistent with the recent history (Methods – Equation 14; **Figure 4D**). Humans and animals are capable of using probabilistic inference to determine the change points in reversal learning paradigms (Behrens, Woolrich, Walton, & Rushworth, 2007; Costa, Tran, Turchi, & Averbeck, 2015). Since inference is a model-based strategy, it is often contrasted with habits. However, we find that combining inference of the high-level context with habitual responses to low-level states allows for maximizing reward in both contexts without sacrificing behavioral flexibility (**Figure 4E**; Supplementary Figure S4B). In other words, animals should overmatch in dynamic environments if they are able to learn the underlying structure. Conversely, they should adopt a conservative matching strategy when they can’t infer this structure.

To test this, we reanalyzed data from two studies that probed monkeys on probabilistic reversal learning under different conditions (Costa et al., 2015; Kubanek & Snyder, 2015). Briefly, monkeys chose between pairs of stimuli through saccades or arm movements and received probabilistic juice reward, and the reward assigned to each stimulus reversed randomly across blocks of trials (Methods). Costa et al.’s experiment employed binary rewards with low variability in block lengths (**Figure 5A** – top), while in Kubanek et al.’s study both reward magnitudes and block lengths were exponentially distributed and difficult to anticipate (**Figure 5A** – bottom). The hierarchical model above predicts that Costa’s monkeys can habitually choose the best option (i.e., overmatch) and still adapt to reversals by inferring switches in context, while Kubanek’s monkeys would make more exploratory choices as the underlying task structure is too complex to enable contextual inference. These predictions are borne out in the data (**Figure 5B** – top vs bottom; Supplementary Figure S5). In further agreement with this, adaptation dynamics in Costa et al. was well captured by a contextual inference process model that switches between two habitual responses (**Figure 5B** – top inset). To more directly test the contribution of goal-directed and habit systems, we fit a regression model that expressed monkeys’ action, *a_t_*, on any given trial *t* as a function of previous trial’s action, *a_t−_*_1_, and outcome, *a_t−_*_1_*r_t−_*_1_, where *a* ∈ {−1, 1} and *r* ∈ {−1, 1} (Methods). The regression weight on the previous action was substantially larger than previous outcome except on trials immediately following a reversal (**Figure 5C** – left) suggesting that Costa’s monkeys tend to perseverate before and after adapting. This result was also readily explained by the model that uses inference to switch between habits (**Figure 5C** – right). A similar regression analysis on Kubanek’s monkeys yielded an opposite pattern of results where the previous outcome was more influential than previous action (**Figure 5D** – left), a finding that was explained by a model in which actions were controlled by goal-directed rather than habit system (**Figure 5D** – right).

**Figure 5:**
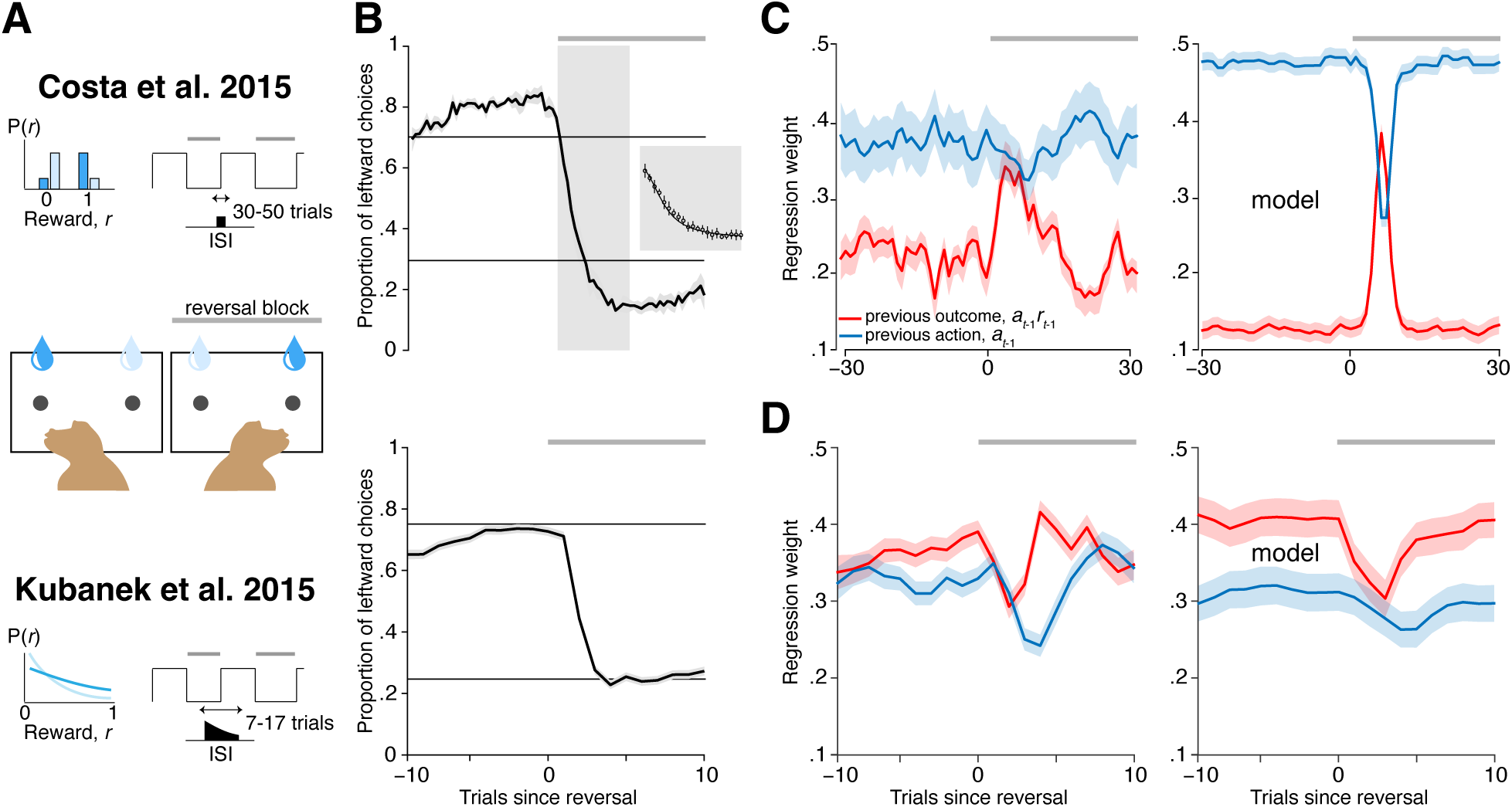
Theory explains behavioral heterogeneity in reversal learning. **A.** Top: Monkeys chose between leftward/rightward targets and obtained a fixed reward with a certain probability (*p*=0.7 for correct target, *p*=0.3 for incorrect). The side of the correct target reversed every 30-50 trials in a predictable manner. Bottom: Similar to (**A**), but in a noisier environment – both the reward ratio of the two targets and the interval between reversals were drawn from truncated exponential distributions (mean reward ratio of 0.75; mean interval of 11 trials). **B.** Top: Monkeys exhibited substantial overmatching, yet successfully adapted to reversals. Horizontal lines correspond to matching law. Inset: Adaptation dynamics in the first 20 trials after reversal (circles) are well explained by a model that combined habits with context inference (line). Bottom: Monkeys in the noisier environment adapted to reversals while exhibiting perfect matching behavior. **C.** Left: The effect of previous trial’s outcome (red) and previous trial’s choice (blue) on the monkey’s choice at various time points before and after the reversal. Right: Similar analysis on data simulated from the model that combined habits with context inference. **D.** Left: Similar to (**C**–left), but for monkeys in the noisier environment. Right: Similar analysis on data simulated from the model driven by the goal-directed system.

### 3.4 Habits in perceptual discrimination

The theory can also offer insights about behavior in settings where the source of uncertainty in outcome is internal rather than external. Concretely, our focus so far has been on experiments that probe behavior using probabilistic rewards and simple stimuli. In this case, the state of the world, *s*, can be unambiguously inferred from the observed stimulus, *o*, and uncertainty arises solely from the probabilistic nature of rewards. In contrast, experiments on perceptual decision-making tend to use noisy stimuli and deterministic rewards. In such cases, the mapping from states to rewards is unambiguous, but instead there is uncertainty in one’s belief about the state, denoted by *P* (*s*|*o*). The above analyses still apply for such tasks since uncertainty in the state propagates to reward, *P* (*r*|*o*) = Σ*_s_P* (*r*|*s*)*P* (*s*|*o*) ∝ *P* (*s*|*o*). Consequently, perceptual reports can be habitized if the probability of choosing an action exceeds the confidence associated with the inferred state (Supplementary Figure S6A). Thus, the theory offers an interpretation for exploratory choices governing ‘lapses’ in perceptual decisions (Pisupati, Chartarifsky-Lynn, Khanal, & Churchland, 2021) as a deliberate strategy to resist habit formation.

A surprising implication emerges in perceptual tasks that require accumulating evidence from time-varying observations *o_t_*. In such tasks, animals sometimes tend to give more weight to observations at earlier time points even when later observations are equally informative about the underlying state *s*. Existing accounts of this primacy effect implicate different aspects of the accumulation process, such as bounded diffusion (Kiani, Hanks, & Shadlen, 2008), competitive inhibition (Tsetsos, Gao, McClelland, & Usher, 2012; Keung, Hagen, & Wilson, 2020) and online feedback of decision-related information to sensory areas (Lange, Chattoraj, Beck, Yates, & Haefner, 2021), the latter of which highlights the impact of decision-making systems on evidence accumulation. Therefore, we hypothesized that the the temporal weighting of evidence could be influenced by the outcome of arbitration between goal-directed and habit systems. To test this, we constructed a model featuring both learning and inference (**Figure 6A**). In this model, the output of the winner-take-all mechanism WTA(**Q***_t_,* **H***_t_*) at any given time step *t*, contributes to evidence accumulation by functioning as a prior for performing belief updates i.e., inference in the subsequent time step *t*+1 (Methods – Equation 15). After viewing all T observations, an action is selected following which **Q** and **H** are themselves updated to enable learning (Supplementary Figure S6B).

**Figure 6:**
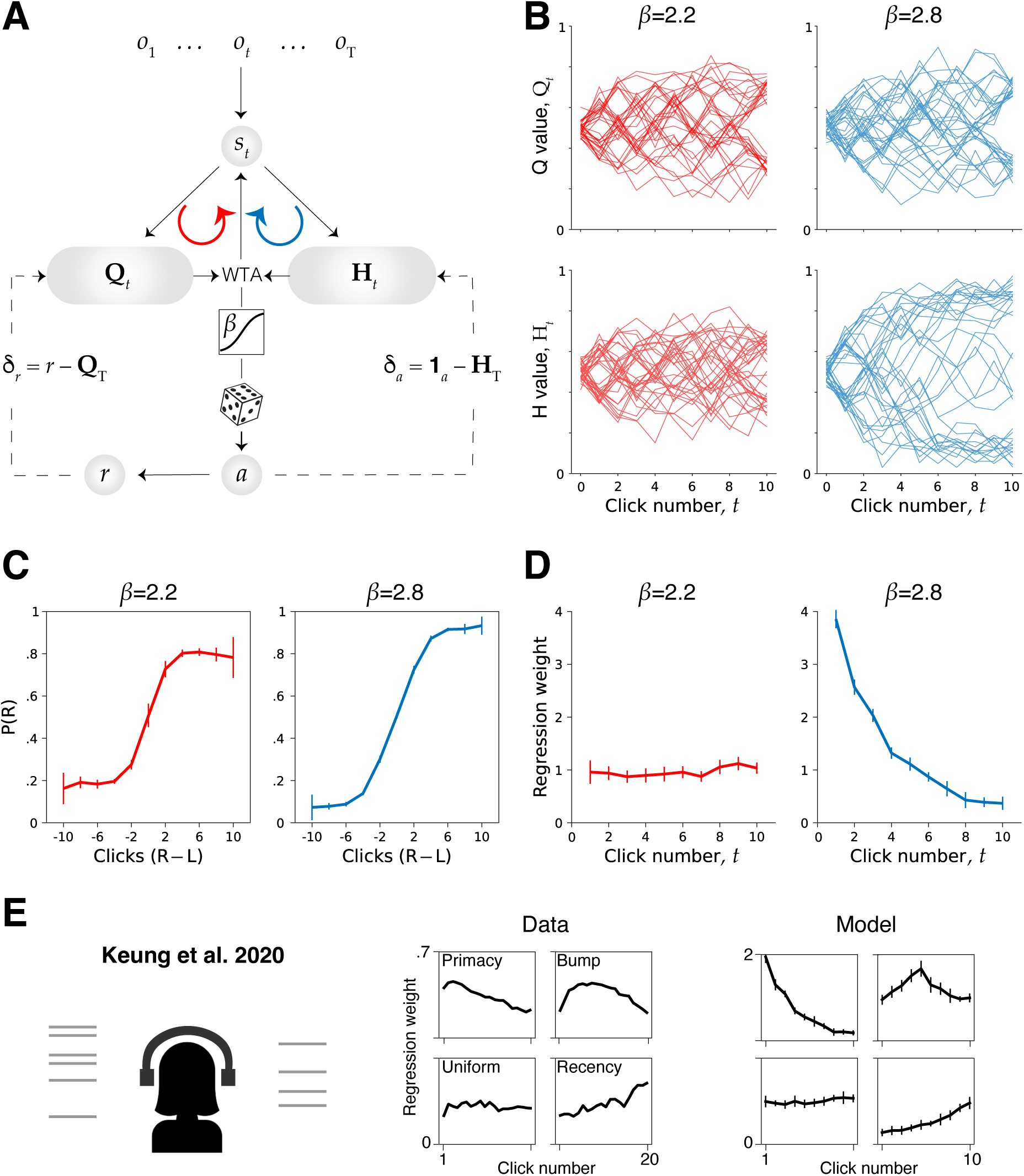
Effect of goal-directed and habit systems on weighting of sequential evidence. **A.** Schematic of the model adapted for learning and inference in the auditory clicks task. **B.** Action value and preference are tied to belief states and evolve during evidence integration. Value and preference dynamics in a subset of trials stimulated from models with low (*β* = 2.2, left) and high (*β* = 2.8, left) degree of exploitation. **C.** Psychometric curves of models with different degrees of exploitation, showing the probability of rightward choice against evidence strength. **D.** Psychophysical kernel of models with different degrees of exploitation, showing the weighting of individual clicks as a function of their position in the sequence. **E.** Heterogeneity in temporal weighting profiles across humans peforming the auditory clicks task (middle) is recapitulated by the model across different regimes (right).

We simulated the model on an auditory clicks task by varying the softmax parameter *β* to tradeoff exploration and exploitation, and analyzed the performance after learning process had converged. On each trial, the model is presented with a sequence of binary observations and asked to report the true underlying state by choosing the corresponding action (Methods). We found that action value **Q***_t_* tracks the true belief across time within each trial, while action preference **H** is underconfident or overconfident when *β* is low (exploratory regime) or high (exploitative regime) respectively (Methods; **Figure 6B**).

Consequently, depending on the regime, belief updates are governed by the goal-directed system or by the habit system, yielding different lapse rates (**Figure 6C**; Supplementary Figure S6C). We characterized the temporal weighting of evidence by regressing the chosen action against observations presented at different time points during the trial (Methods). In the exploratory regime, the model weights all observations uniformly since feedback from the goal-directed system accurately tracks the true belief state, enabling optimal evidence accumulation (**Figure 6D – left**). In contrast, the model exhibits primacy effect i.e., overweights early observations in the exploitative regime (**Figure 6D – right**). This is because the habit system makes an overconfident determination of the belief state based on initial evidence, and feedback from this system weakens the contribution of subsequent evidence by biasing their interpretation towards the initial belief. In addition to uniform weighting and primacy effect, other forms of temporal weighting profiles have been observed in the auditory clicks task including bump and recency effects (Keung et al., 2020), and variants of the model recapitulated this heterogeneity in weighting profile (**Figure 6E**). Specifically, recency effects emerge when the integration process underlying belief updates is leaky as suggested in previous works, while bumps in weighting profile are seen to arise when the source of decision-related feedback switches from goal-directed to habit system in the middle of this leaky integration process. This latter result raises the possibility that arbitration between goal-directed and habit systems might be more dynamic than commonly assumed.

### 3.5 Theory explains striatal manipulation effects on behavior

Our theoretical results are based on the assumption that learning in the goal-directed and habit systems are mediated by different types of prediction errors. Previous works have attributed goal-directed and habitual behaviors to dorsomedial (DMS) and dorsolateral (DLS) striatum respectively (Yin et al., 2004, 2005; Gremel & Costa, 2013). Since plasticity in the striatum is thought to depend on prediction errors signaled by dopamine (Bromberg-Martin, Matsumoto, & Hikosaka, 2010), we asked whether dopaminergic inputs to DMS and DLS are consistent with reward prediction errors and action prediction errors respectively. A recent study examined the effects of stimulating dopamine terminals in these areas throughout operant training (Seiler et al., 2022). In this paradigm, mice learned to nosepoke to obtain a scheduled reward (**Figure 7A** – top). Critically, dopamine stimulation (in DMS or DLS) coincided either with rewarded or random nosepokes in separate groups of mice, testing the extent to which the effects of stimulation, if any, are reward-dependent (**Figure 7A** – bottom). When we simulate learning in this paradigm with a relatively high degree of exploitation (Methods), we find two qualitatively different learning regimes – an early, reward-dependent regime in which learning is mediated by the goal-directed system, and a late, reward-independent regime in which learning is mediated by the habit system (**Figure 7B**).

**Figure 7:**
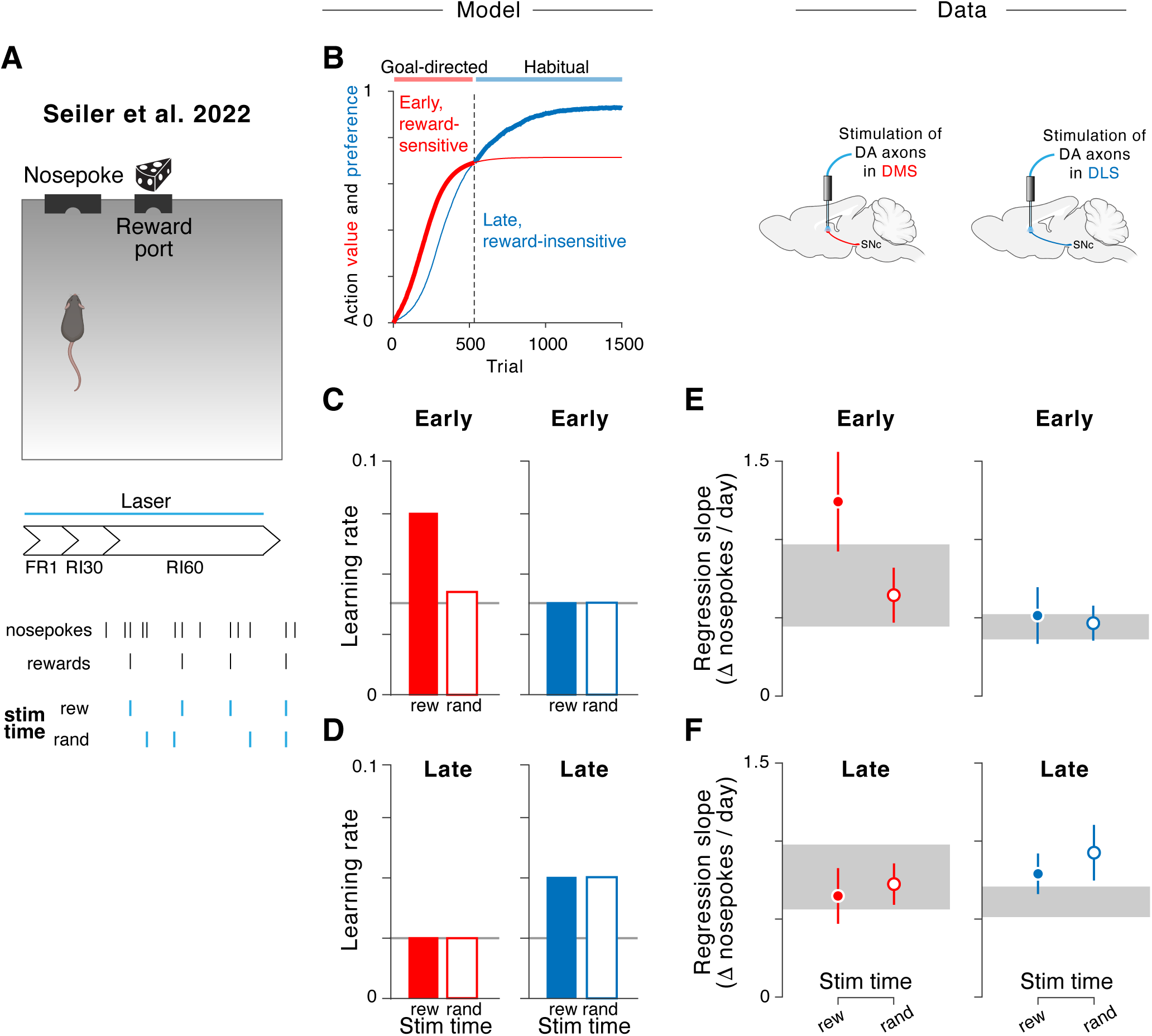
Dopamine stimulation effects are consistent with distinct prediction errors in DMS and DLS. **A.** Mice nosepoked to obtain scheduled rewards that got progressively more sparse with training, while dopamine axon terminals in DMS or DLS were optogenetically stimulated at time points of rewarded nosepokes (‘rew’) or random nosepokes (‘rand’). Adapted from (Seiler et al., 2022). **B.** Simulated learning curves for relative value, *Q* (red) and preference, *H* (blue) for lever press actions show an early, reward-sensitive learning phase (*Q > H*) followed by a late, habitual learning phase (*H > Q*). **C.** Simulated effects of ‘rew’ and ‘rand’ stimulations applied to the reward prediction error (left) and action prediction error (right) signals on the rate of learning in the goal-directed regime. **D.** Similar to (**C**) but for effects in the habit regime. **E.** Effects of dopamine stimulation in DMS (left) and DLS (right) during early learning (FR1+RI30). **F.** Similar to (**E**), but during the late learning (RI60) phase. Gray line in **C-D** denote the baseline learning rate in the absence of stimulation. In **E-F**, gray shaded regions denote 95% CI of learning rates in the control condition and error bars denote 95% CI. FR1: fixed ratio-1, RR30: random ratio-30, RR60: random ratio-60, DMS: dorsomedial striatum, DLS: dorsolateral striatum.

We first simulated the effects of dopamine stimulation in the model separately in the goal-directed and habit systems, assuming that stimulation amplifies prediction error signals in these systems (Methods). We found that stimulation in the goal-directed system was effective in enhancing the learning rate in the early phase when learning was mediated by this system (*Q > H*), but not later (**Figure 7C,D** – left panels). Moreover, since learning in this system is mediated by reward prediction errors, the increase was appreciable only when stimulation was restricted to rewarded actions (‘rew’ vs ‘rand’). In contrast, stimulation in the habit system enhanced the learning rate in the late learning phase (*Q < H*) but not earlier (**Figure 7C,D** – right panels). Critically, because action prediction errors are not influenced by reward, the strength of this effect was comparable for stimulation of rewarded and random actions.

If learning in the goal-directed and habit systems is mediated by reward and action prediction errors respectively, the effects of dopamine stimulation in DMS and DLS should parallel the above simulation results. To test this, we partitioned the mouse data into early and late training sessions, and quantified the rate at which level press actions increased across sessions within each phase (Methods). In alignment with model predictions, stimulating DMS dopamine substantially increased the learning rate in the early training phase but not later, and specifically when the stimulations were yolked to rewarded nosepokes (**Figure 7E,F** – left panels; Supplementary Figure S7A). On the other hand, stimulating DLS dopamine did not have any effect in the early phase, but yielded a modest increase in learning rates in the late phase even when stimulations targeted random (unrewarded) nosepokes (**Figure 7E,F** – right panels; Supplementary Figure S7B). These results suggest that dopamine signals in DMS and DLS mediate qualitatively different forms of learning.

We demonstrated earlier that the theory can be used to understand behavior in paradigms not explicitly designed to probe habits such as evidence accumulation. To investigate whether this extends to the interpretation of circuit function, we considered a recent study by (Bolkan et al., 2022). In this study, the authors examined the effect of optogenetically silencing DMS neurons in mice trained to perform evidence accumulation in a virtual T-maze (**Figure 8A**). They found that silencing DMS neurons affects behavior when cues may be present on either side but not in a task variant lacking distractor cues (**Figure 8B** – top vs bottom), suggesting that engagement of DMS depends on cognitive demands. To understand why this might be, recall that when the source of uncertainty is internal like in this task, probability matching requires making choices with a probability that is equal to the confidence in the inferred state while overmatching causes decisions to become habitized. Because confidence i.e., posterior belief is, on average, much lower in the more cognitively demanding task variant, mice must reduce their degree of exploitation (*β*) in order to maintain good performance (Supplementary Figure S8A). Therefore, we simulated the model separately to learn the two task variants, allowing greater exploitation in the variant with no distractors (Methods). The models developed different strategies, relying on the goal-directed system to perform the task that required evidence accumulation and the habit system to perform the task with no distrators (Supplementary Figure S8B). Consequently, inhibiting the goal-directed system selectively impaired the performance in the cognitively demanding task and spares the easy task (**Figure 8C** – red), recapitulating the effects of DMS silencing in mice. Interestingly, inhibiting the habit system yielded the opposite pattern of results (**Figure 8C** – blue). Since DLS is thought to be engaged in habitual behaviors, we predict that silencing DLS should selectively impair performance in the task with reduced cognitive demands.

**Figure 8:**
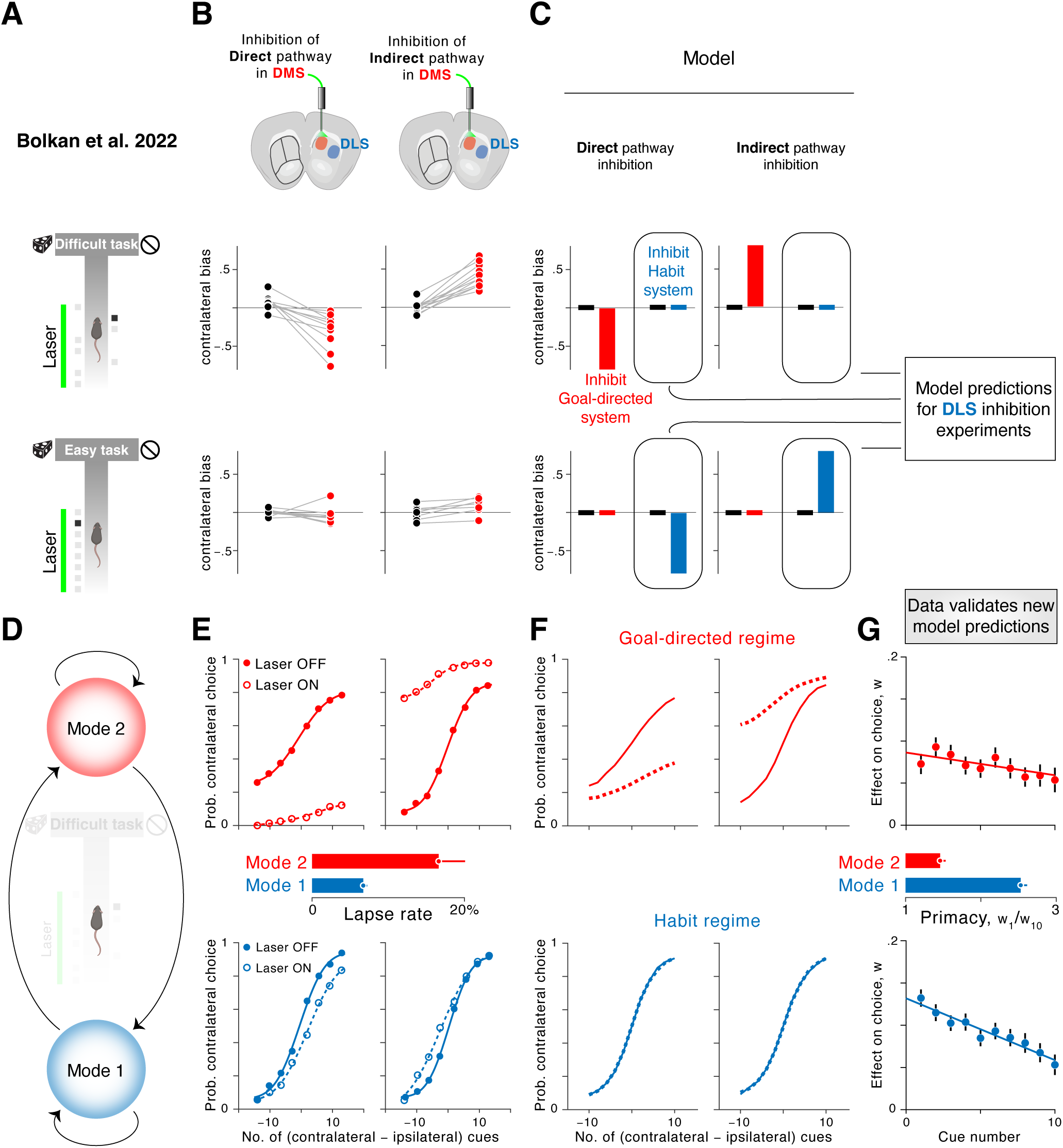
DMS inhibition reveals signatures of alternation between goal-directed and habit regimes. **A.** Top: Mice navigated a virtual T-maze, turning leftward or rightward depending on the side of the corridor that had more cues (difficult task). Laser stimulation was applied to inactivate DMS neurons during the cue integration period. Bottom: In a separate variant of the task, cues were only present on one of the two sides on any given trial and required no integration (easy task). Adapted from (Bolkan et al., 2022). **B.** Left: Behavioral effects of inactivating the direct pathway DMS neurons in the two task variants. Right: Similar to left, but showing effects of inactivating the indirect pathway DMS neurons. Black dots denote the baseline performance in the absence of inhibition. **C.** Simulated effects of inhibiting either the goal-directed system (red) or the habit system (blue) on performance in the two task variants. Black bars denote baseline performance. **D.** Unsupervised methods identify different latent behavioral modes in mice during the cue integration task. **E.** Psychometric curves (filled circles), lapse rates (horizontal bars), and effects of DMS inactivation (open circles) in the two behavioral modes. **F.** Simulated effects of inhibiting the goal-directed system on the psychometric curves when the model performance is governed by the goal-directed (top) and habit system (bottom). **G.** The psychophysical kernel of mice, showing the temporal weighting profile of cues in the two behavioral modes. Horizontal bars show the primacy effect, quantified as the ratio of the weighting between the first and the last cue. Error bars denote 95% CI.

At a more granular level, (Bolkan et al., 2022) fit a latent variable model and found that mice alternated between different behavioral modes even within the difficult task variant. Mice paid little to no attention to sensory cues in one of these modes, so we focus on the other two modes showing high level of task engagement (**Figure 8D**). Mice exhibited more exploratory behavior in mode 2 than in mode 1 (**Figure 8E** – red vs blue solid lines), resulting in significantly higher lapse rates (**Figure 8E** – middle). Since exploitation promotes habitization, the theory predicts that modes 1 and 2 are likely under the control of the habit system and goal-directed system respectively. Consistent with this, DMS silencing affected behavioral performance only in mode 2 (**Figure 8E** – red vs blue dashed lines). These results are readily reproduced by simulating the model in two regimes that differ solely in their degree of exploitation (**Figure 8F**). To more directly test whether modes 1 and 2 correspond to habitual and goal-directed behavioral modes respectively, we analyzed the temporal weighting of different sensory cues by mice i.e., the psychophysical kernel within each behavioral mode. We showed in the previous section that goal-directed and habit systems exert distinct effects on the temporal weighting profile in evidence accumulation, with the latter system causing a primacy effect in which early cues are weighted more (Figure 6D). We found that mice indeed signficantly overweighted early cues in mode 1 than in mode 2 (**Figure 8G**; Supplementary Figure S8C), suggesting that the differential effect of DMS inhibition across behavioral modes is due to an alternation between goal-directed (exploration) and habitual (exploitation) behavioral strategies.

## 4 Discussion

Most learned behaviors are initially goal-directed. When do they transform into habits? We presented a mathematical theory tracing the emergence of habits to the trade-off between exploration and exploitation. It identifies probability matching as a natural reference point to determine if a behavioral policy is exploiting (overmatching) or exploring, yielding the principle that exploitation leads to habit formation while exploration preserves goal-directed behavior. While this theoretical result likely represents an idealization which stems from the assumed winner-take-all arbitration mechanism, we demonstrated that it can explain a broad range of qualitative phenomena at the behavioral and neural levels, and make testable predictions.

### 4.1 Habit as a learning trap

The result might seem counterintuitive at first glance because habits are characterized by insensitivity to rewards, while exploitation indicates a high affinity to rewards. However, this view only considers habit expression. The theory, on the other hand, reflects the influence of exploration-exploitation trade-off on the learning process underlying habit formation. From a learning perspective, the results become intuitive: in noisy environments, exploitation causes stimulus-action covariance to gradually exceed action-reward covariance, thereby masking the latter. This harks back to the idea linking habits to correlation-based law of effect (Baum, 1973; Dickinson, 1985) but our model expands its scope by pitting it in competition against a parallel learning system that implements Thorndike’s law of exercise following recent works (Miller et al., 2019; Bogacz, 2020). The resulting model enabled us to identify a clear demarcation point for habit formation in the form of probability matching (not to be confused with Herrnstein’s matching law; see (Mongillo et al., 2014) for review) which requires a precise balance between exploration and exploitation to ensure that goal-directed and habit systems achieve equal performance. Behavior will remain goal-directed as long as exploitation stays at or below this critical level, explaining why extended training does not lead to habits in some cases (DeRusso et al., 2010; Robbins & Costa, 2017; Garr et al., 2021; LaFlamme et al., 2022). Otherwise, behavior converges to a stable equilibrium point where action preferences become independent of the outcome, falling into a habit. This view of habits is conceptually similar to learning traps that arise from limited exploration in multi-alternative decision-making settings, but differs in two ways. First, previous models attribute learning traps to knowledge blind spots created by incomplete information sampling such as hot stove and hot hand effects (Rich & Gureckis, 2018). In contrast, our model shows that when two systems compete, one system (habit) can get trapped even if the other system (goal-directed) has perfect knowledge. Second, learning traps result in suboptimal performance (Teodorescu & Erev, 2014; Raab & Hartley, 2020) whereas habits maximize rewards and become detrimental only later if action values change. Nonetheless, habits may be viewed as a learning trap in that they arise from an interplay between learning and exploitative decision-making.

### 4.2 Relation to prior work

A large body of work is based on the influential idea that goal-directed and habitual behaviors can be understood as model-based (MB) and model-free (MF) strategies respectively (Daw et al., 2005; Dolan & Dayan, 2013). MB strategies prospectively evaluate the consequences of taking actions, while MF strategies rely on previously learned stimulus-action associations. A widely adopted two-step task used to distinguish between these strategies revealed that reduced MB strategy is correlated with habits, evaluated traditionally by devaluation sensitivity (Gillan et al., 2015). However, reduced MB control might reflect an inability to form perfect models rather than increased MF control (Collins & Cockburn, 2020; da Silva & Hare, 2020; Castro-Rodrigues et al., 2022). Moreover, since MF learning is mediated by reward prediction errors, this strategy cannot explain persistent lever-pressing under manipulations that induce negative reward prediction errors e.g., during contingency reversals (Dezfouli & Balleine, 2012) and superimposed omissions (Dickinson, Squire, Varga, & Smith, 1998). To overcome these limitations, we adopt a slightly different view that maps goal-directed and habitual behaviors to value-based (VB) and value-free (VF) strategies respectively (Miller et al., 2018, 2019). This view of habits retains the computational efficiency of MF algorithms while making it impenetrable to rewards. Because we focused on single-step decisions in this study, we implemented the goal-directed system also as a MF system but we note that the theory is agnostic to how action values are computed. MB planning methods (Simon & Daw, 2011; Mattar & Daw, 2018; Ho et al., 2022; Lakshminarasimhan et al., 2023) can replace or augment the MF implementation of goal-directed system in the current model. In fact, to simulate outcome devaluation here (Figure 3), we assumed that offline consumption of rewards instantly alters action values which would require MB computations. Thus, while most studies focus on *how* goal-directed and habit systems compute – MB or MF, the proposed theory is based on *what* these systems compute – VB or VF. This shift in emphasis coupled with the assumed winner-take-all (WTA) arbitration mechanism, allows for the two systems to be in conflict even after prolonged experience, leading to qualitatively different regimes of behavior compared to models featuring cooperative interaction like summation (Perez & Dickinson, 2020). Though this may seem idealized, it opens the door for the theory to be applied beyond the confines of the operational definition of habits. We demonstrated this by applying it to reversal learning and evidence accumulation, identifying a shared mechanism for disparate phenomena like overmatching, choice history effect, primacy bias etc. As a concrete example, our finding that habits cause confirmation bias might explain why choice history biases the drift rate of accumulation and not just the starting point (Urai et al., 2019). This finding also offered new insight into data from (Keung et al., 2020) and (Bolkan et al., 2022), suggesting that evidence accumulation is subject to dynamic shifts between habitual and goal-directed control at fast (within trial) and slow (across trials) timescales. Similar shifts have been reported in rats performing a two-step decision making task (Venditto et al., 2024), and constitute an important avenue for future research on how competition between decision-making systems shapes the dynamics of cognition.

Another key aspect of this study is examining how the representations on which learning mechanisms operate impact habit formation. In particular, we showed that learning on abstract state representations informed by hierarchical inference enables one to switch between habits in response to probabilistic reversals in lieu of supplanting the original policy. The theory correctly predicted data showing that overmatching in probabilistic reversal learning is seen only in settings where context inference is possible (Costa et al., 2015; Kubanek & Snyder, 2015) and broadly consistent with the finding that habits are more context-sensitive than goal-directed actions in operant conditioning (Thrailkill & Bouton, 2015). This is also in alignment with previous work showing that learning on extended state representations can enable model-free agents to masquerade as model-based (Akam et al., 2015) and reminiscent of a parallel but distinct idea of habits as hierarchically organized action sequences (Dezfouli & Balleine, 2012). The latter study proposes that when model-based computations are applied on action chunks, they produce policies in which individual actions appear habitual, implying that habits belie resource-rational computations (Nassar et al., 2018; Lai et al., 2022). Future studies should look into how both state and action representations jointly influence habit formation.

### 4.3 Neurobiology of habits

The theory applies to neural systems that learn to choose actions in parallel using distinct types of prediction error signals. Convergent lines of evidence from experiments on rodents suggest an anatomical basis for this model. First, causal experiments have mapped habitual and goal-directed actions to dorsolateral (DLS) and dorsomedial striatum (DMS) (Yin et al., 2004, 2005), areas homologous to the primate putamen and caudate respectively. Second, plasticity in these regions is modulated by dopamine (Fisher et al., 2017) and dopamine depletion impairs both behavioral strategies (Faure, Haberland, Condé, & Massioui, 2005; Lex & Hauber, 2010). Third, the two regions receive inputs from different sets of dopaminergic neurons in the substantia nigra that have qualitatively different response properties (Lerner et al., 2015; Tsutsui-Kimura et al., 2020). Fourth, transsynaptic tracing studies reveal a parallel circuit architecture with minimal crosstalk between the two nigrostriatal circuits (Ambrosi & Lerner, 2022).

For learning mechanisms in this circuit to be aligned with the model, dopamine signals in DMS and DLS must convey reward prediction errors (RPE) and action prediction errors (APE) respectively. While there is evidence for RPE signals in DMS (Parker et al., 2016), dopamine signaling in the DLS has proved challenging to interpret. A recent study in which mice self-stimulated dopamine release in the DMS or DLS by pressing one of two levers found that only the DMS group adapted when the stimulating lever was switched, suggesting that DLS dopamine promotes habits (van der Merwe et al., 2023). Our analyses of data from (Seiler et al., 2022) complements the above finding by showing that dopamine stimulation in DMS and DLS during operant learning produces behavioral effects akin to theoretical RPE and APE signals, respectively. Our conclusions are based on partitioning the data into early and late training phases, and more work is needed to test the robustness of these findings by tracking dopamine signals throughout the learning process. Interestingly, (Greenstreet et al., 2022) performed such analyses in an auditory discrimination paradigm and found signatures of APE signals in the striatal tail and propose this as a locus of habit formation. It is possible that APE signals are diffuse throughout multiple striatal regions with different regions performing value-free learning on different sensory representations, analogous to feature-specific RPEs (Lee et al., 2024).

Another neural constraint is that the competition between goal-directed and habit systems is resolved downstream, meaning that both systems are active even if only one of them controls behavior. This might explain why neural activity in DMS and DLS evolves independently in parallel (Ito & Doya, 2015) and remains correlated with behavior even after extensive training (Vandaele et al., 2019). Consistent with this, behavioral and pharmacological manipulations can unmask habitual and goal-directed strategies (Coutureau & Killcross, 2003; Hardwick et al., 2019), and synaptic changes associated with the initial task acquisition are not erased after contingency reversal (Ghosh & Zador, 2021). However, other studies suggest an asymmetric reorganization of activity in DMS to DLS via interference (Yin et al., 2009; Thorn et al., 2010; Bergstrom et al., 2018; Turner et al., 2022). Notably, the above studies all employed paradigms that required substantial movement and thus an element of motor skill learning. Since skills and habits are not exactly the same (Robbins & Costa, 2017), it is possible that the former involves interference between circuits whereas habit formation does not.

### 4.4 Assumptions and extensions

We made several simplifying assumptions that future experimental validation and extensions of this work must consider. First, we assume a constant level of random exploration throughout learning (fixed *β* parameter), but humans sometimes use more sophisticated forms of exploration (Schulz et al., 2020; Fox et al., 2023). Second, since the policy mapping function controlling exploration operates downstream of the winner-take-all arbitration mechanism, the theory cannot account for the rapid switching between strategies suggested by our analysis of (Bolkan et al., 2022). Moreover, this precludes the use of algorithms that explicitly learn a policy such as policy-gradient and actor-critic methods that enjoy experimental support (Bennett et al., 2021; Coddington et al., 2023). Above limitations can be tackled by funneling the output of both systems through separate softmax operators with their own *β*, before arbitrating between them. Finally, we have not explored the emergence of stereotypy in naturalistic behaviors with continuous state and action spaces (Markowitz et al., 2018; Lakshminarasimhan et al., 2018, 2020; Stavropoulos et al., 2022; Alefantis et al., 2022; Zhu et al., 2022). Developing a biologically plausible account of habits in such settings might require rethinking the learning signals and state representations (Lindsey & Litwin-Kumar, 2022; K. Yamada & Toda, 2023). Approaches like meta-learning are especially useful in this regard (Jiang & Litwin-Kumar, 2021; Shervani-Tabar & Rosenbaum, 2023; Lakshminarasimhan et al., 2024).

### 4.5 Conclusion

Despite a substantial body of empirical work, the fundamental principles governing habit formation have remained elusive. We propose a principle linking habit formation to the trade-off between exploration and exploitation. This expresses habits in more general terms, paving the way both for using a wider range of experimental paradigms as well as leveraging inter- and intra-individual variability in exploratory behavior to investigate the underlying mechanisms.

## 5 Methods

### 5.1 Model

Let *s*, *a*, and *r* denote state, action, and reward respectively. The model receives state *s* as input, chooses action *a* as output, and obtains reward *r* as feedback for learning. The model comprises two systems. The goal-directed system learns to estimate the value, *Q_s,a_*, i.e., average reward associated with each state-action pair. The habit system learns to estimate the tendency, *H_s,a_*, to choose a particular action in a given state. We refer to these estimates as *action value* and *action preference* respectively, following the conventional names given to estimates learned by value-based and value-free methods (Sutton & Barto, 2018). States, actions, and rewards are all binary unless specified otherwise. Below, we describe the learning rule for updating action values and action preferences, the rule for arbitrating between them, and the decision rule for choosing an action.

***Learning rule.*** The model learns through trial and error to iteratively improve estimates *Q_s,a_*(*k*) and

*H_s,a_*(*k*) where *k* denotes the trial number. Learning in the goal-directed system is given by:

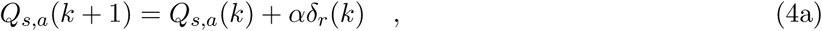

where *δ_r_*(*k*) = *δ_aak_ r* +(1 − *δ_aak_*) (1 − *r*) − *Q_s,a_*(*k*) denotes an action-specific *reward prediction error* (RPE) and *α* is the learning rate. Here, *δ_aak_* is the Kronecker delta function which evaluates to 1 for the action *a^k^* chosen on trial *k* i.e., *a* = *a^k^*, and 0 for the non-chosen action i.e., *a* ≠ *a^k^*. This rule allows for updating action values associated with both the chosen and non-chosen actions assuming that taking the non-chosen action would have produced the opposite outcome, 1 − *r*, and yields asymptotically accurate estimates provided reward probabilities associated with the two actions sum to 1. For learning tasks where the latter condition is not met, we only update the value associated with the chosen action. Simultaneously updating both action values is not essential, but mirrors learning in the habit system where both action preferences must be simultaneously updated to ensure that learning is unbiased.

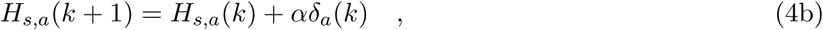

where *δ_a_*(*k*) = *δ_aak_* − *H_s,a_*(*k*) denotes the *action prediction error* (APE). Note that it is imperative to update the action preference associated with the non-chosen action for learning in this system to converge to the relative frequency of taking actions. Otherwise, both action preferences will approach 1.

***Arbitration rule.*** In any given state *s*, we arbitrate between the two systems by applying a greedy winner-take-all (WTA) rule that compares the vector of action values and action preferences associated with the current state, *Q_s_* and *H_s_*. When rewards are binary and the reward probabilities associated with the two actions sum to 1 as assumed above, both *Q_s_* and *H_s_* will have similar scaling since ∥*Q_s_*∥_1_ = ∥*H_s_*∥_1_ = 1 where ∥ · ∥ _1_ denotes the 1-norm. In this case, the WTA rule simply chooses the vector whose element has the largest magnitude as follows:

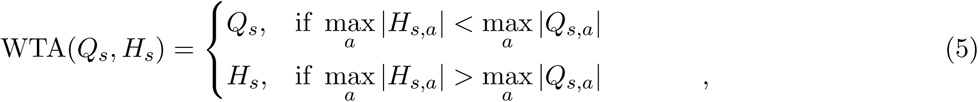

where ties are broken randomly. In case reward probabilities associated with the actions do not sum to 1 or if rewards have different magnitudes, then *Q_s_* and *H_s_* will have different scaling. In such cases, we divide *Q_s_* by ∥*Q_s_*∥_1_ to ensure that ∥*Q_s_*∥_1_ = 1 before applying the WTA rule. The division transforms absolute values into relative values, as observed in the monkey brain (Padoa-Schioppa, 2009; H. Yamada, Louie, Tymula, & Glimcher, 2018). This rescaling was used to simulate operant conditioning tasks.

***Decision rule.*** The estimates of the system selected by the WTA rule are transformed into a probability distribution by applying a policy mapping function. We use the softmax operator, which is parametrized by the inverse-temperature parameter *β* which controls the trade-off between exploration and exploitation. The probability of choosing action *a* is given by:

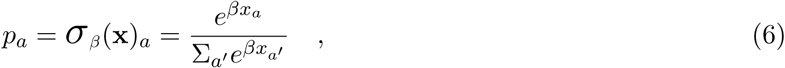

where **x** = WTA(*Q_s_, H_s_*) denotes the output of the arbitration rule described above. For binary actions, the chosen action is given by *a* ∼ Bernoulli(*p_a_*). Note that the probability of receiving a reward is 1 − *p_r_* ≤ *p_a_p_r_* + (1 − *p_a_*)(1 − *p_r_*) ≤ *p_r_*, where *p_a_* is the probability of taking an action and *p_r_* is the reward probability associated with that action.

### 5.2 Asymptotic Action Value and Action Preference

To calculate the asymptotic action values, we take Equation 4a to the limit of convergence:

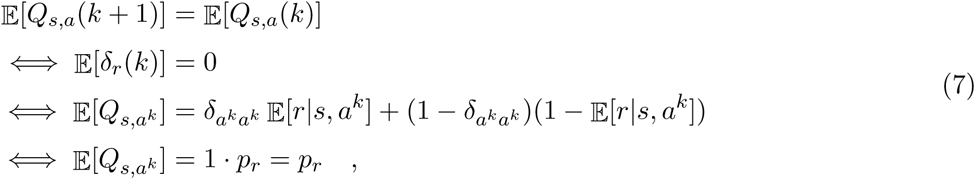

where E[·] denotes the expectation over outcomes and we have assumed, without loss of generality, that the probability of outcome *r* = 1 is *p_r_* and *r* = 0 is 1 − *p_r_* when taking action *a^k^* in state *s*. Likewise, substituting *a* = *a^\k^* in line 3 above, we get E[*Q_s,a\k_*] = 1 − *p_r_*. Thus, *Q_s,a_* converges to the true reward probability corresponding to action *a* in state *s*. To calculate the asymptotic action preferences, we take Equation 4b to the limit of convergence:

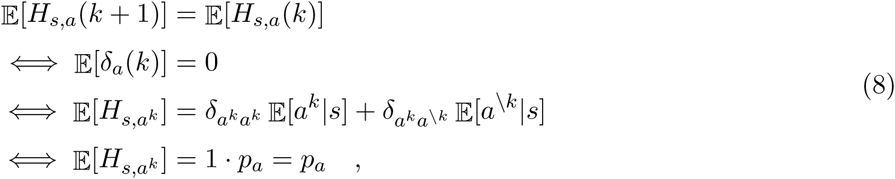

where E[·] is the expectation over actions and we have assumed, without loss of generality, that the probability of taking action *a^k^* is *p_a_*. Since the probability of taking the other action *a^\k^* is 1 − *p_a_*, we get E[*H_s,a\k_*] = 1 − *p_a_*. Thus, *H_s,a_* converges to the true probability of taking action *a* in state *s*.

### 5.3 Critical level of exploitation for habit formation

Let *β*_crit_ denote the critical level of exploitation needed to promote habit formation. The value of *β*_crit_ that induces habits depends on the task structure. We derive the expression for *β*_crit_ for some special cases below by making use of the fact that actions are governed by the goal-directed system when *β < β*_crit_ and the two systems are matched when *β* = *β*_crit_.

***Symmetric, binary rewards.*** Suppose *r* = 1 with probability *p_r_* and *r* = 0 with probability 1 − *p_r_* for one action, and vice-versa for the other. In this case, *Q_s_* = (*p_r_,* 1 − *p_r_*) and *H_s_* = (*p_a_,* 1 − *p_a_*) from Equations 7 and 8 respectively. We then have:

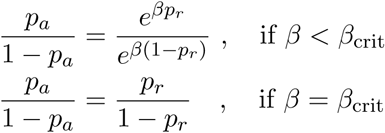

Combining the above equations, we get:

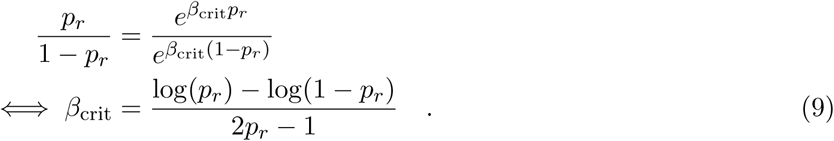

***Asymmetric and/or arbitrary rewards.*** Suppose *r* = *r*_1_ with probability 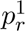 and *r* = 0 with probability 1 − 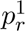 for action *a* = *a*_1_, and *r* = *r*_2_ with probability 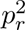 and *r* = 0 with probability 1 − 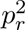 for *a* = *a*_2_. Adapting the derivation of Equation 7 to this case, the estimates of the goal-directed system will converge to the expected value, 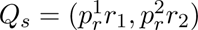. The estimates of the habit system derived in Equation 8 do not depend explicitly on the reward statistics, *H_s_* = (*p_a_,* 1 − *p_a_*). As we mentioned earlier, whenever *Q_s_* and *H_s_* have different scales, we rescale elements of *Q_s_* to express them as relative value:

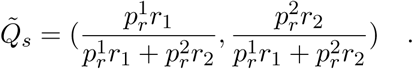

Following the steps used to derive Equation 9, we get:

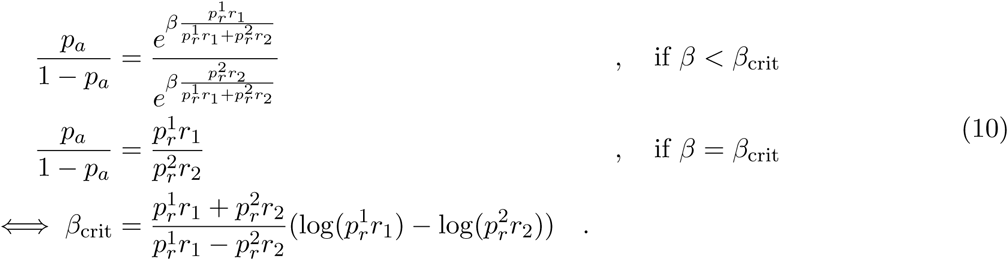

***Free-operant conditioning.*** In this setting, the animal is free to perform an action e.g., lever press and receives a reward *r* delivered according to a predetermined schedule e.g., variable ratio or variable interval. This is unlike tasks with a well-defined trial-structure, in which each trial corresponds to some state *s*. To apply the theory to this setting, we divide time into bins and reset the (subjective) timer whenever a reward is obtained, such that each bin corresponds to a unique state. In each bin, we assume that the animal can either perform an action with (a state-dependent) probability *p_a_* to harvest reward *r* with (a state-dependent) probability *p* or relax and receive a unit reward with probability 1 such that 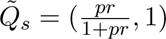. Following the steps used to derive Equation 9, we get:

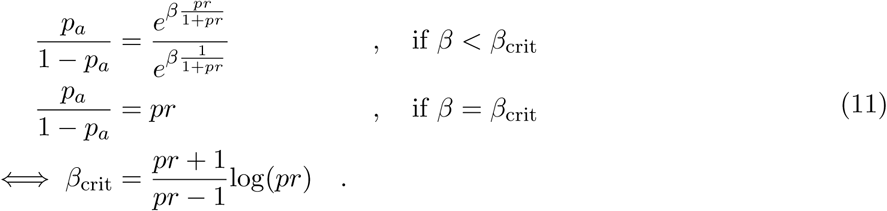

***Action sequences.*** Suppose the task requires a sequence of two actions (e.g., chain pull and lever press) to be performed in order to obtain reward *r*. Let *a*^D^ and *a*^P^ denote the distal and proximal actions, and let 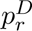 and 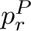 denote the probability that those actions culminate in a reward. If the probability of successfully completing a sequence is *η*, then 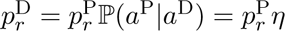. Substituting in Equation 11 yields the minimum *β* that habitizes the distal action.

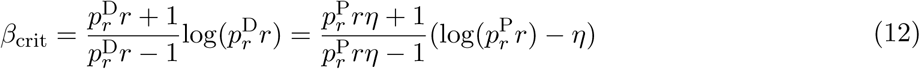

***Delayed rewards.*** Consider a variant of the free-operant conditioning paradigm discussed above in which actions lead to reward *r* with probability *p* after a time delay *T*. To capture the effect of this delay on RPE, we multiply it by an eligibility trace 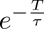 which decays exponentially with some time constant *τ*. This reduces the asymptotic action value estimated by the goal-directed system to 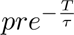 but spares the habit system. Substituting in Equation 9 yields the minimum *β* for habit formation in the delayed reward setting.

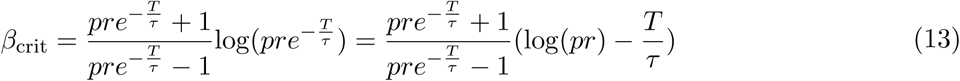

### 5.4 Simulation details

We simulated the model using a learning rate *α* = 0.01 in both the goal-directed and habit systems. In all cases, we ran simulations of the model with 10 different seeds of the pseudorandom generator and report their average. For plots showing theoretical estimates as a function of task parameters, estimates were calculated by binning parameters into approximately 50 bins. Task-specific details are described below.

***Basic learning task.*** For simulations of the basic learning task in Figure 1, states, actions and rewards were all binary. The probability of reward for choosing the best and worst action in a given state was *p_r_*= 0.8 and 1−*p_r_*= 0.2 respectively, and the level of exploitation was either *β* = 1.5 or *β* = 3. The action value and action preferences were initialized to 0.5 i.e., *H_s,a_*(0) = *Q_s,a_*(0) = 0.5 ∀ *s, a*. Because of the symmetric structure of the task, action values and preferences of both the chosen and non-chosen action were updated at the end of each trial. Simulations were run for 500 trials. For the theoretical estimates plotted in Figure 2, we calculated each estimate for a range of parameters *p_r_* ∈ [0, 1] and *β* ∈ [1, 4].

***Free-operant tasks.*** For the action sequence task in Figure 3A, the distal action yielded a reward of *r* = 15 when followed by the proximal action and no reward otherwise. Furthermore, we assumed *r* = 1 when forgoing an opportunity to take an action to reflect savings in metabolic cost. For plots showing theoretical estimates, we varied the probability, P(*a*^P^|*a*^D^) = *η*, of taking a distal action followed by a proximal action i.e., success rate in the range *η* ∈ [0.25, 1] and *β* ∈ [1, 4]. For simulations to qualitatively match data from (Balleine et al., 1995), we roughly estimated the success rate in the behavioral data to be *η* = 0.9 from the ‘actions per opportunity’ plot in the original study, and used *β* = 2.8. For the delayed reward task in Figure 3B, we assumed *r* = 15 when a lever press leads to a reward and *r* = 0 otherwise. As before, *r* = 1 when not pressing the lever. The time constant of the eligibility trace was *τ* = 10*s*. For the theory plots, we varied the delay *T* between lever press and reward in the range *T* ∈ [0, 30*s*] and *β* ∈ [1, 4]. For simulations to qualitatively match data from (Urcelay & Jonkman, 2019), we used a delay of *T* = 20*s* from the original study, and *β* = 2.8. For comparing reward schedules in Figure 3C, we simulated fixed interval (FI60) and variable interval (VI60) tasks. We divided time since last reward into bins of 1s, and estimated action value and preference for each time bin. We assumed *r* = 100 when a lever press leads to a reward and *r* = 0 otherwise, and as before, *r* = 1 when not choosing not to press the lever. We simulated different levels of uncertainty in the reward schedule by interpolating between FI60 and VI60 using gamma distributed schedules with a fixed mean of 60s and varying standard deviations in the range *σ* ∈ [0, 60*s*]. For simulations to qualitatively match data from (DeRusso et al., 2010), we used FI60 and VI60 schedules with *β* = 2.8. Because the rate of lever pressing is initially low in free-operant tasks, in all the above simulations, we initialized *H_a_*(0) = *Q_a_*(0) = 1 for *a* = 0 i.e., choosing no action, and *H_a_*(0) = *Q_a_*(0) = 0 for *a* = 1 i.e., lever pressing. After learning, we tested the model under devaluation by scaling down the action value for lever press in both schedules by a factor of 2.

***Reversal learning.*** The task structure of the simulations in Figure 4 was similar to the basic learning task, except the reward contingency was switched after 250 trials. This meant that if taking action *a* in state *s* was rewarded with probability *p_r_*, the reward probability for that state-action pair switched to 1 − *p_r_* after reversal. Similar to the simulation of the basic learning task, the reward probability was *p_r_* = 0.8 and the level of exploitation was either *β* = 1.5 or *β* = 3. To infer a switch in the context *c*, we use Bayes’ rule:

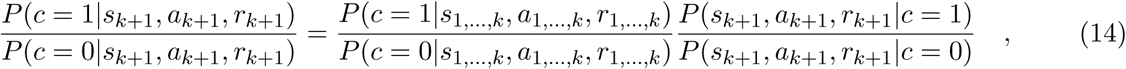

where *P* (**c**|*s, a, r*) denotes the posterior probability distribution over contexts, *P* (*s, a, r*|**c**) denotes the likelihood of the contexts based on the observed (*s, a, r*), and subscripts denote trial indices. Note that for the task we simulated, *P* (*s, a, r* = 1|*c* = 1) is 0.8 if (*s* = 0*, a* = 0) or (*s* = 1*, a* = 1), and 0.2 otherwise, and vice-versa for *P* (*s, a, r* = 0|*c* = 1) as well as *P* (*s, a, r* = 1|*c* = 0). The ratio of the posteriors is clipped to 1000. For simulations in Figure 4E where context inference was used in conjunction with learning in the goal-directed and habit systems, the action values and preferences depended both on state *s* and context *c*, and were initialized to *H_c,s,a_*(0) = *Q_c,s,a_*(0) = 0.5 ∀ *c, s, a*. We used a relatively high degree of exploitation, *β* = 3 to demonstrate the benefit of using context inference in conjunction with habits.

***Evidence accumulation.*** Simulations in Figure 6 correspond to a decision making task in which evidence from a sequence of noisy observations *o_t_* (clicks) drawn from one of two states *s* (left or right) must be accumulated. A deterministic reward *r* = 1 is delivered for choosing the action *a* that correctly identifies the underlying state. Although the true state is binary, the belief state, defined as the posterior probability over the state at any given time *t*, is continuous-valued between 0 and 1. Therefore, we binned the belief into 7 uniformly spaced bins and learned action values *Q_b,a_* and action preferences *H_b,a_* for each discrete belief state denoted by *b* ∈ {1*, …,* 7}. Although a choice is not made until after the final time step, feedback from one of the two systems affects the evidence accumulation process according to Bayes’ rule where the feedback at each time step is taken to be the prior for the subsequent time step:

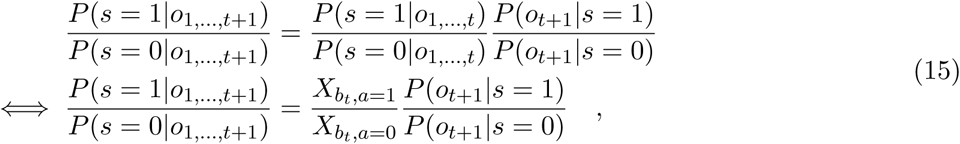

where *X_bt_* = WTA(*Q_bt_, H_bt_*) is the output of the winner-take-all rule and *b_t_* is the discretized belief at time *t*. We simulated the task for sequences of length 10 with *β* set to either 2.2 or 2.8 to obtain psychophysical kernels that exhibited no bias or primacy bias respectively. To qualitatively reproduce kernels from (Keung et al., 2020), we simulated two more variants. To simulate recency bias, we used *β* = 2.2 and introduced a leak constant of 0.8 that shifted the posterior ratio towards 1. Simulations to obtain nonmonotonic kernels also employed leaky integration but the level of exploitation increased from *β* = 2.2 to *β* = 3.5 after the first 5 samples of the sequence.

***Dopamine stimulation.*** The basic simulation in Figure 7 was similar to the description above for free-operant tasks with *r* = 2.5 and *β* = 3. The experiments by (Seiler et al., 2022) used progressively sparse reward schedule, but in the simulations we used a VI60 schedule throughout learning for simplicity. Over the course of learning, the model transitioned from the goal-directed regime to habit regime. We used this transition point to delineate early and late phases of learning and analyzed the effect of stimulation in each phase separately. The precise mechanism by which dopamine stimulation affects learning is not known. For the simulations, we assume that stimulation of dopamine axon terminals in the striatum reinforces actions by amplifying positive prediction errors by a factor of 2 while leaving negative prediction errors unchanged, *δ*_stim_ = *k* · *δ* where *k* = 2 ·]_*_δ>_*_0_. This means that dopamine stimulation amplifies long term potentiation in the direct pathway neurons and/or long term depression in the indirect pathway neurons that are active during lever pressing. We considered and ruled out alternatives that were inconsistent with key aspects of the data. For example, amplifying both positive and negative prediction errors in the model resulted in no net effect on behavior, while modeling stimulation by adding a non-negative constant to the prediction error led to similar predictions for scrambled (‘rand’) and rewarded (‘rew’) stimulations.

***Striatal inhibition.*** For simulations in Figure 8, we used methods similar to the one described above for evidence accumulation. As mentioned in the main text, overexploitation deteriorates performance in the difficult version of the task where cues are presented on both sides (left and right) in the same trial but not in the version that lacks distractor cues in panel 8C. Therefore, we used *β* = 2.2 and *β* = 3.5 to simulate the difficult and easy variants respectively. Each trial of the original experiment contained up to 20 cues, but we used sequences of length 10 in our simulations for simplicity. Because the DMS inhibition experiments in (Bolkan et al., 2022) targeted direct and indirect pathway neurons separately, we adapted the model by decomposing the action value and action preference estimates as follows:

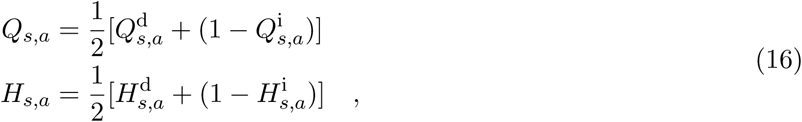

where superscripts d and i denote estimates in the direct and indirect pathways respectively. Learning rule in the direct pathway was identical to Equation 4, while that for the indirect pathway was similar except the identity of the action *a^k^* chosen on a given trial indexed by *k* was flipped by setting 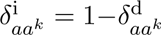 before applying the learning rule. For simulating unilateral, pathway-specific inhibition of a striatal region, we downscaled the estimate corresponding to the contralateral action (0 or 1) in the corresponding pathway (d or i) of the corresponding system (*Q* or *H*) by a factor of 5. The qualitative findings were robust to the precise factor used in the simulations. For panel 8F, we simulated two separate models with different levels of exploitation that solved the difficult task variant in either the goal-directed (*β* = 2.2) or habitual (*β* = 2.8) regimes.

### 5.5 Data acquisition

The data panels in Figure 3A, 3B, 3C and 6E summarizing results of previous studies were obtained by feeding an image of the original figure panels from (Balleine et al., 1995), (Urcelay & Jonkman, 2019), (DeRusso et al., 2010), and (Keung et al., 2020) respectively through WebPlotDigitizer v5.1 (Rohatgi, 2024), a computer vision assisted software for data extraction, occasionally intervening to select the data points manually. This is somewhat imprecise but we deem this sufficient because the simulations are only meant to capture qualitative aspects of the data. In all other cases, we used the data files provided by the authors of the original study. We used publicly available data from (Costa et al., 2015) and (Kubanek & Snyder, 2015) for the analyses of reversal learning behavior in monkeys shown in Figure 5. Data for DMS/DLS dopamine stimulation experiments in Figure 7 and DMS inhibition experiments in Figure 8 were provided by (Seiler et al., 2022) and (Bolkan et al., 2022) respectively.

### 5.6 Data analysis

Our re-analyses of data from previous studies involved fitting simple regression models to assess the influence of different predictors on behavior. The predictors were chosen to test specific hypotheses described below.

***Reversal learning.*** To test whether behavior deviated from matching law, we first reproduced the analyses performed in the original studies by calculating the proportion of best choices by aligning data from all the experimental blocks to the reversal trial. In the (Costa et al., 2015) dataset (555 blocks, 80 trials/block), we focused on the condition in which the reward probability associated with the actions had a ratio of 0.7:0.3 such that the action probability for matching was 0.7. In the (Kubanek & Snyder, 2015) dataset (1015 blocks, 11 ± 5 trials/block), we analyzed the condition in which the ratio of reward magnitudes was, on average, 3:1 such that the matching action probability was 0.75. To test the hypothesis that overmatching stems from habit formation, we fit a logistic regression model to predict the action *a_k_* chosen in trial *k* from the previous action *a_k−_*_1_ and previous outcome *a_k−_*_1_*r_k−_*_1_ according to:

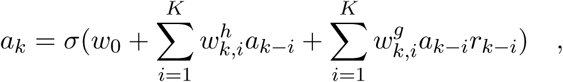

where *σ*(·) is the standard logistic function, *w*_0_ captures a history-independent response bias, and *a, r* ∈ −1, 1. The sums of the absolute value of the regression coefficients 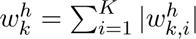 and 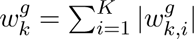 were taken to reflect the influence of the habitual system and the goal-directed system respectively on trial *k* indexed relative to reversal. Due to differences in the block length, we used *K* = 5 and *K* = 1 for (Costa et al., 2015) and (Kubanek & Snyder, 2015) datasets respectively but using *K* = 1 in both datasets yielded qualitatively similar results.

***Dopamine stimulation.*** To test the hypothesis that dopamine stimulation in the DMS and DLS should have different effects during early and late training, we partitioned the data from (Seiler et al., 2022) into two phases. We grouped the training sessions that employed Fixed ratio (FR1) and Random Interval 30 (RI30) schedules together into the ‘early’ phase and subsequent sessions that employed RI60 schedule into the ‘late’ phase. This delineation is imprecise because we do not know if and when behavior switches from goal-directed to habitual mode. In fact, the original study demonstrated a continued involvement of the DMS in punishment resistant reward-seeking using shock probes sessions which we excluded from our analysis. However, the recruitment of DMS after extended training could be context-dependent and thus does not preclude a switch to habitual nosepoking in the non-shock probe sessions following extended training. To test our hypothesis, we fit a piecewise (or segmented) linear regression model to predict the rate of nosepokes as a function of session number across the two training phases. We fit separate models for animals in the ‘control’ group (with no dopamine stimulation), the ‘rew’ group (in which dopamine stimulation coincided with rewarded nosepokes), and the ‘rand’ group (in which dopamine stimulation conincided with random nosepokes), and balanced the datasets from different groups to ensure each group (of 13 ± 1 mice) had an equal number of sessions within each phase. The regression model is defined as:

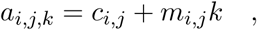

where *i* ∈ {‘control’, ‘rew’, ‘rand’} denotes the group, *j* ∈ {‘early’, ‘late’} denotes the training phase, *k* ∈ {1*, …, k*_max_} denotes the session number, *a* denotes the rate of nosepoking, *c* is the intercept, and *m* denotes the slope of the learning trajectory i.e., learning rate. To ensure continuity between the two phases of piecewise regression, we enforced the constraint *c_i,_*_late_ = *c_i,_*_early_ + *m_i,_*_early_*k*_max_ where *k*_max_ is the number of sessions in the early phase.

***VR navigation.*** The study by (Bolkan et al., 2022) fit a GLM-HMM model to mouse behavior and found three modes. Mice were largely disengaged from the task in mode 3 so we excluded this from our analysis. Notably, they found that unilateral DMS inhibition biased behavior in mode 2 but not in mode 1. We tested the hypothesis that modes 1 and 2 correspond to habitual (driven by DLS) and goal-directed (driven by DMS) regimes respectively. Our simulations of an evidence accumulation model with decision feedback revealed that behavior in the habitual but not the goal-directed regime exhibits primacy bias i.e., a tendency to overweight sensory cues observed early in the sequence. Therefore, we fit psychophysical kernels separately to the set of trials from the two modes, combining the data from all mice (27,257 and 23,756 trials in modes 1 and 2 respectively):

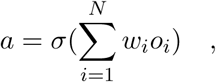

where *a* ∈ {0, 1} denotes the binary choice, and *w_i_* is the influence of the *i*^th^ observation in the sequence *o_i_* ∈ {−1, 1}.

## 6 Code availability

The code for running model simulations is available at https://github.com/kaushik-l/habit-theory.

## 7 Acknowledgements

This work was supported by the Kavli Foundation, the Gatsby Charitable Foundation GAT3780, and a NARSAD Young Investigator Grant #31985 from the Brain & Behavior Research Foundation. We thank Scott Bolkan for sharing the mouse striatal inhibition data, Talia Lerner and Jillian Seiler for sharing the mouse striatal dopamine stimulation data, and the authors involved in (Costa et al., 2015; Kubanek & Snyder, 2015) for sharing their monkey behavioral data. We also thank Ashok Litwin-Kumar, Guillermo Horga, Jean-Paul Noel and Scott Bolkan for providing feedback on this manuscript.

