## Supplemental Figures for "A computational principle of habit formation"

Kaushik Lakshminarasimhan
Zuckerman Mind Brain Behavior Institute, Columbia University, New York, NY, USA

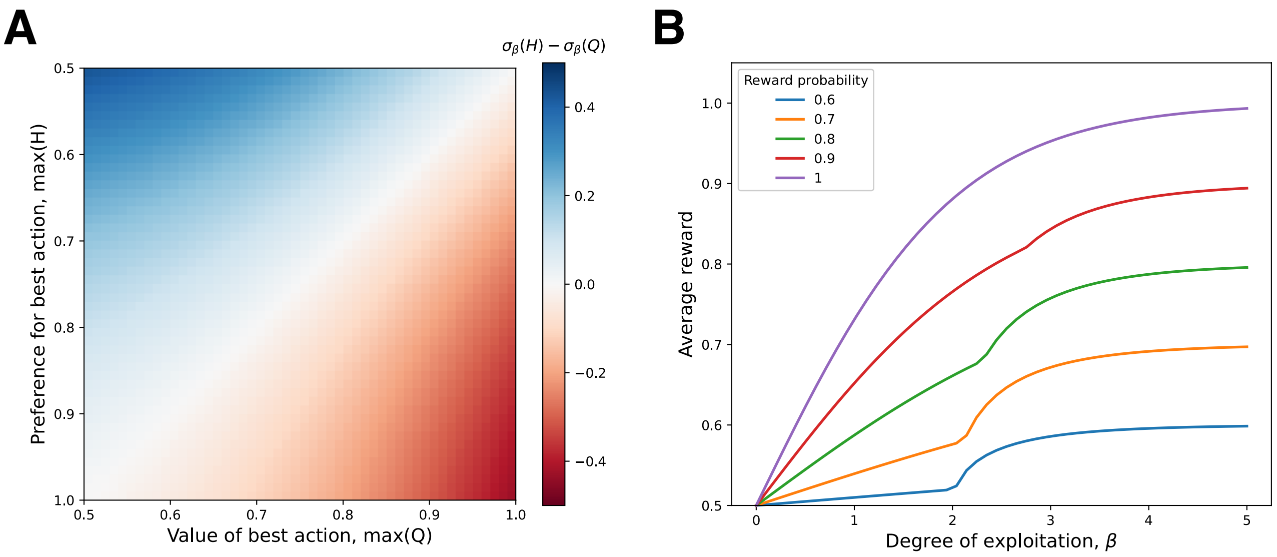


Figure S1: **Arbitration between goal-directed and habit systems**. **A.** The difference between the probability of choosing the best action under habit and goal-directed systems, as a function of the best action's value and the best action's preference. **B.** Average reward harvested by the model as a function of the degree of exploitation, across tasks with different reward probabilities. The kink in the curves correspond to the point at which control shifts from the goal-directed to the habitual system.

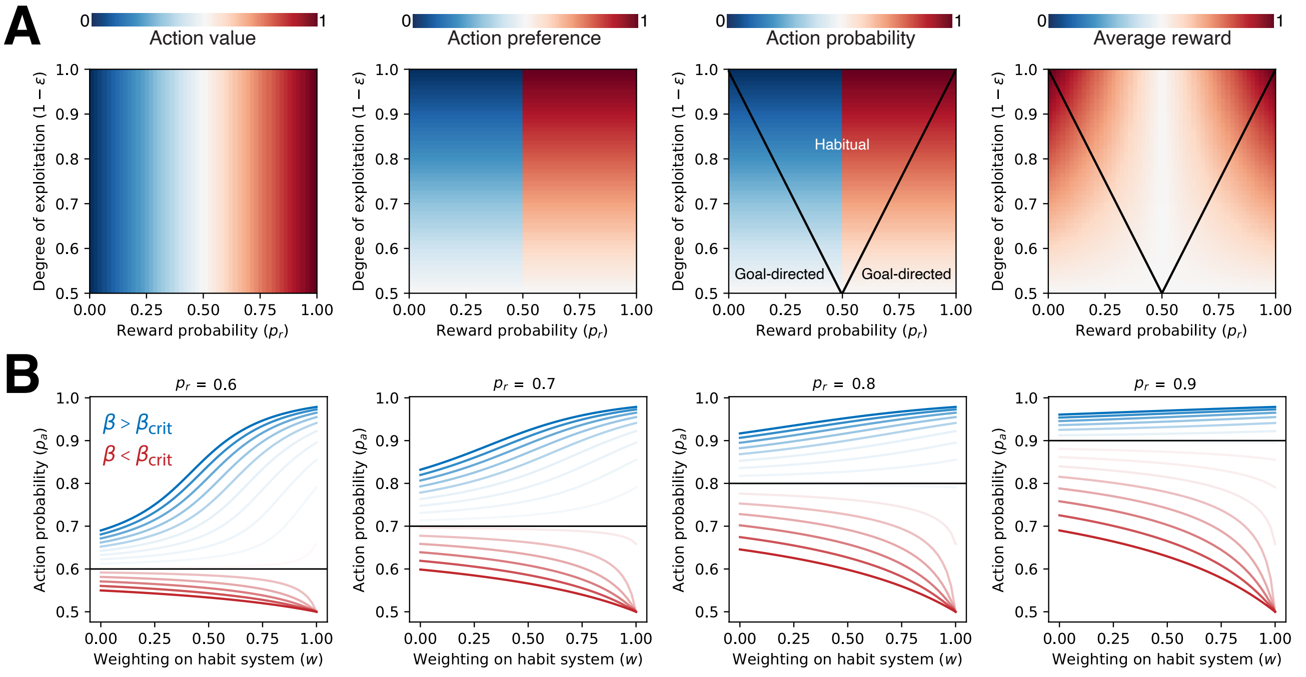


Figure S2: **Performance of model variants**. **A.** Model with $\epsilon$-greedy operator. Action value ($Q$, left), action preference ($H$, left middle), action probability ($p_a$, right middle), and average reward $\bar{r}$ (right) as a function of reward probability $p_r$ and the degree of exploitation $1-\epsilon$ in steady state. Black curve corresponds to the critical level of exploitation $1-\epsilon_{\text{crit}}$ corresponding to matching behavior ($p_a = p_r$). **B.** Model with soft arbitration. The probability of choosing the best action by models in which actions are selected based on a weighted sum of action value $(1-w)$ and action preference $(w)$. Each curve corresponds to a different level of exploitation, ranging from subcritical (dark red) to supracritical (dark blue). Different panels correspond to tasks in which the best action yields different reward probabilities and the black line corresponds to probability matching behavior.

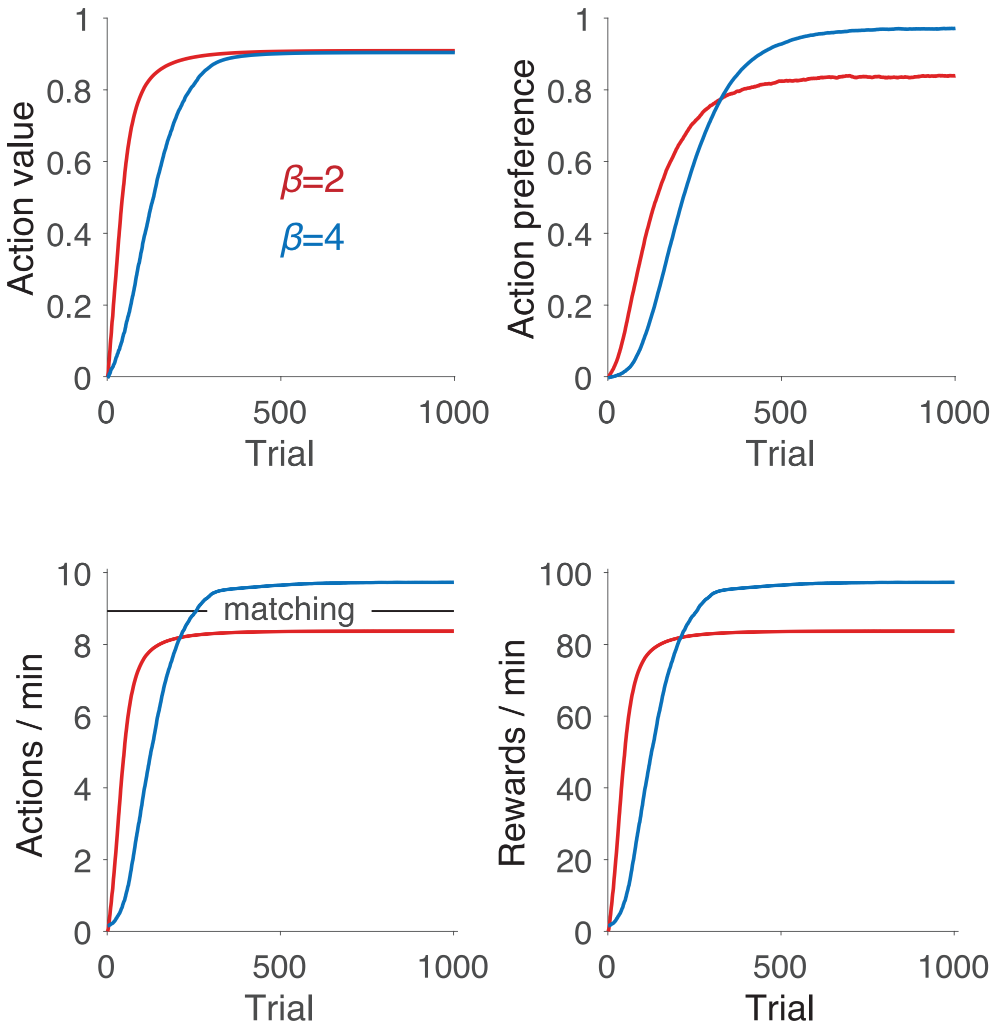


Figure S3: **Modeling free-operant behaviors**. **A.** Action value ($Q$, top left), action preference ($H$, top right), action rate ($p_a$, bottom left), and reward rate $\overline{r}$ (bottom right) in a free-operant (random ratio) paradigm. Red and blue curves correspond to model simulations under low and high degrees of exploitation ($\beta$) respectively. Black horizontal line corresponds to the matching action rate where the fraction of time bins with lever presses is equal to the relative value of the outcomes (of lever pressing and resting).

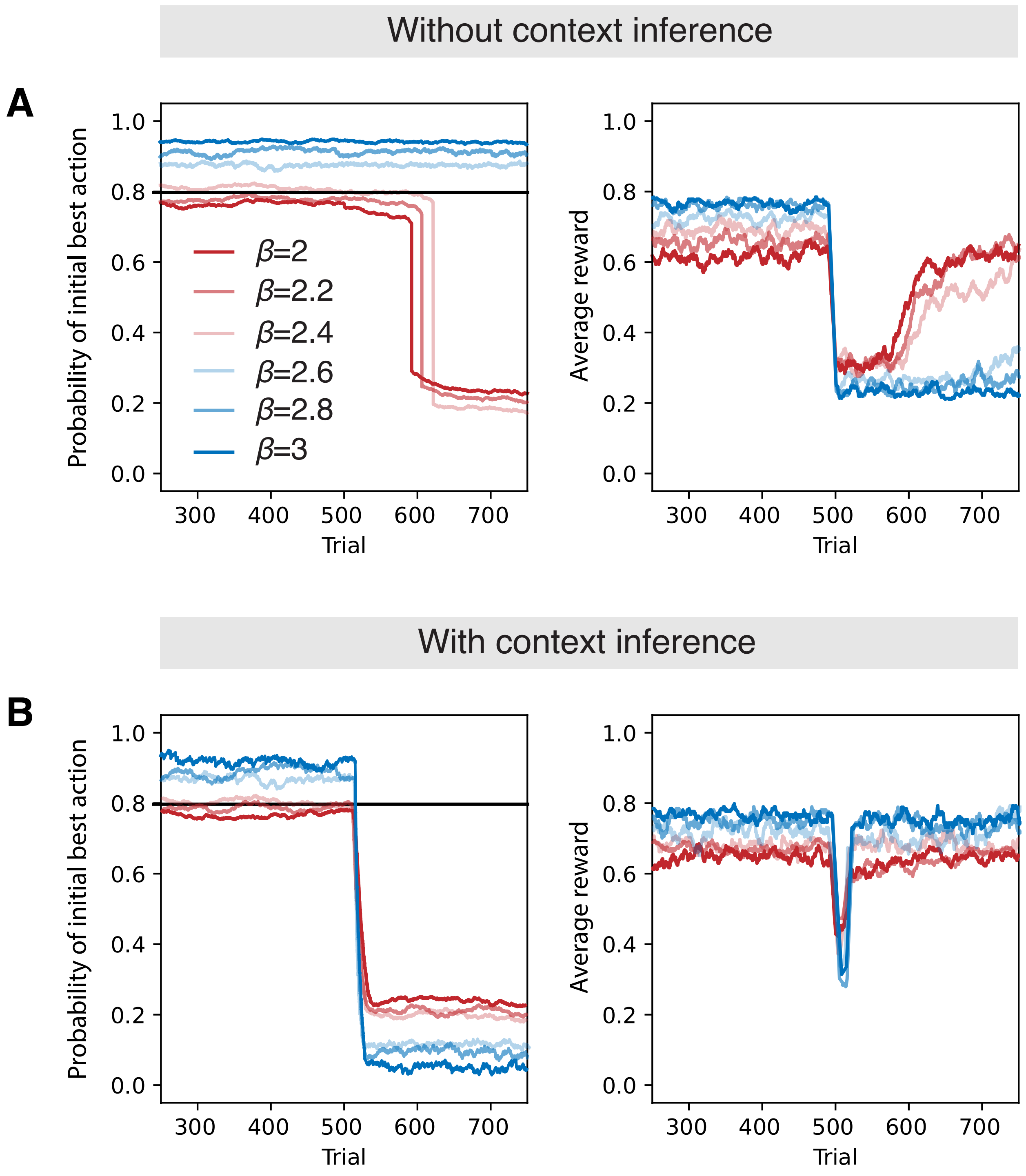


Figure S4: **Reversal learning with and without context inference**. **A.** The probability of taking the initial (i.e., before reversal) best action (left) and the average reward harvested (right) as a function of trials around the reversal point, in a model unaware of context. **B.** Similar to panel **A**, but with probabilistic inference of context. Reversal point corresponds to 500 trials. Each curve corresponds to a different level of exploitation, ranging from subcritical (dark red) to supracritical (dark blue).

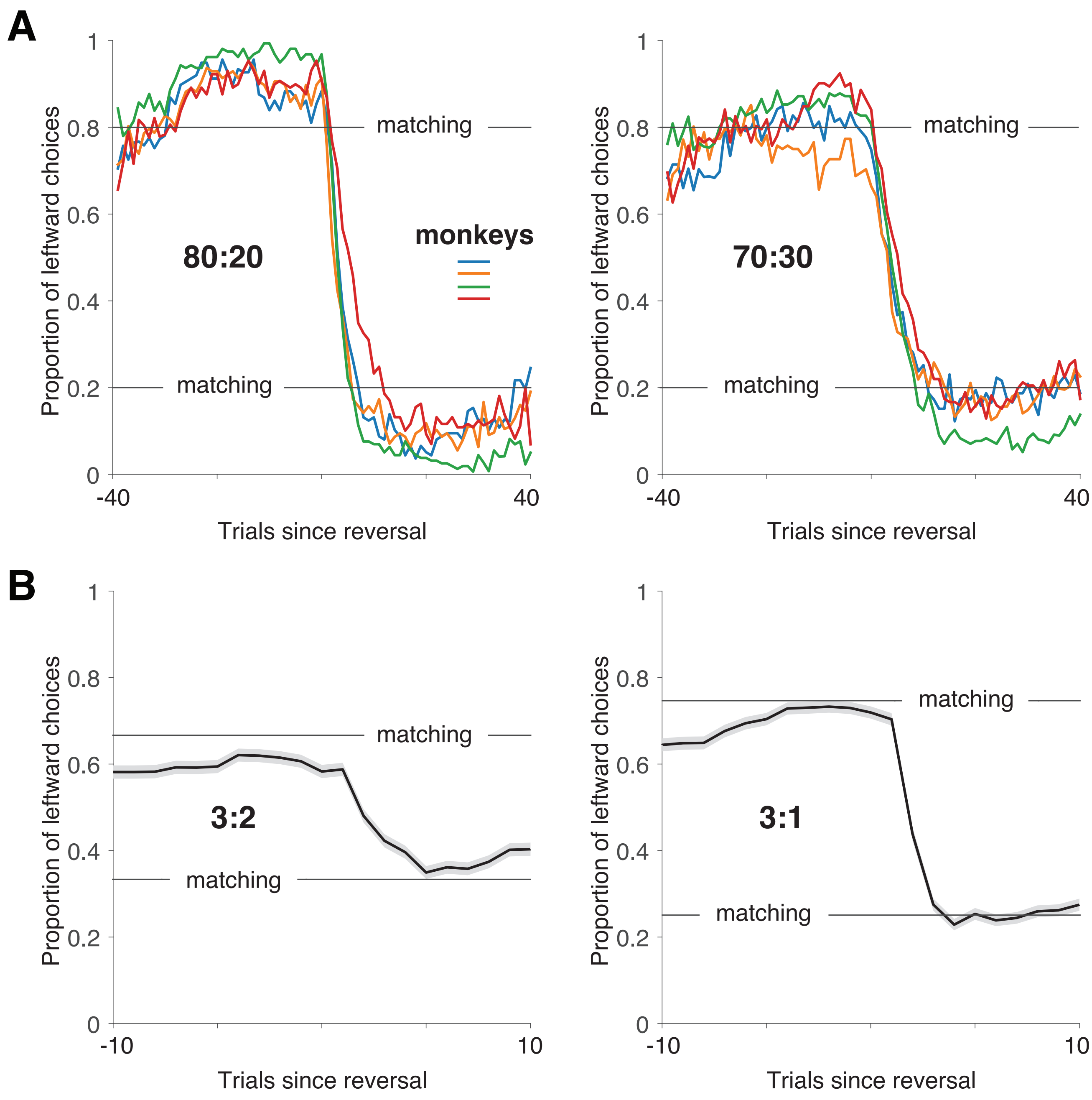


Figure S5: **Reversal learning in monkeys**. **A.** Individual monkeys in the (Costa et al., 2015) dataset exhibited substantial overmatching in both the 80:20 and 70:30 conditions, yet successfully adapted to reversals. Horizontal lines correspond to matching law. In this paradigm, the ratio is expressed as the fraction of trials in which the best action was rewarded. **B.** Monkeys in the (Kubanek & Snyder, 2015) dataset also adapted to reversals but did not overmatch in the 3:2 nor in the 3:1 conditions. In this paradigm, the ratio reflects the relative amount of juice reward dispensed for each action, on average.

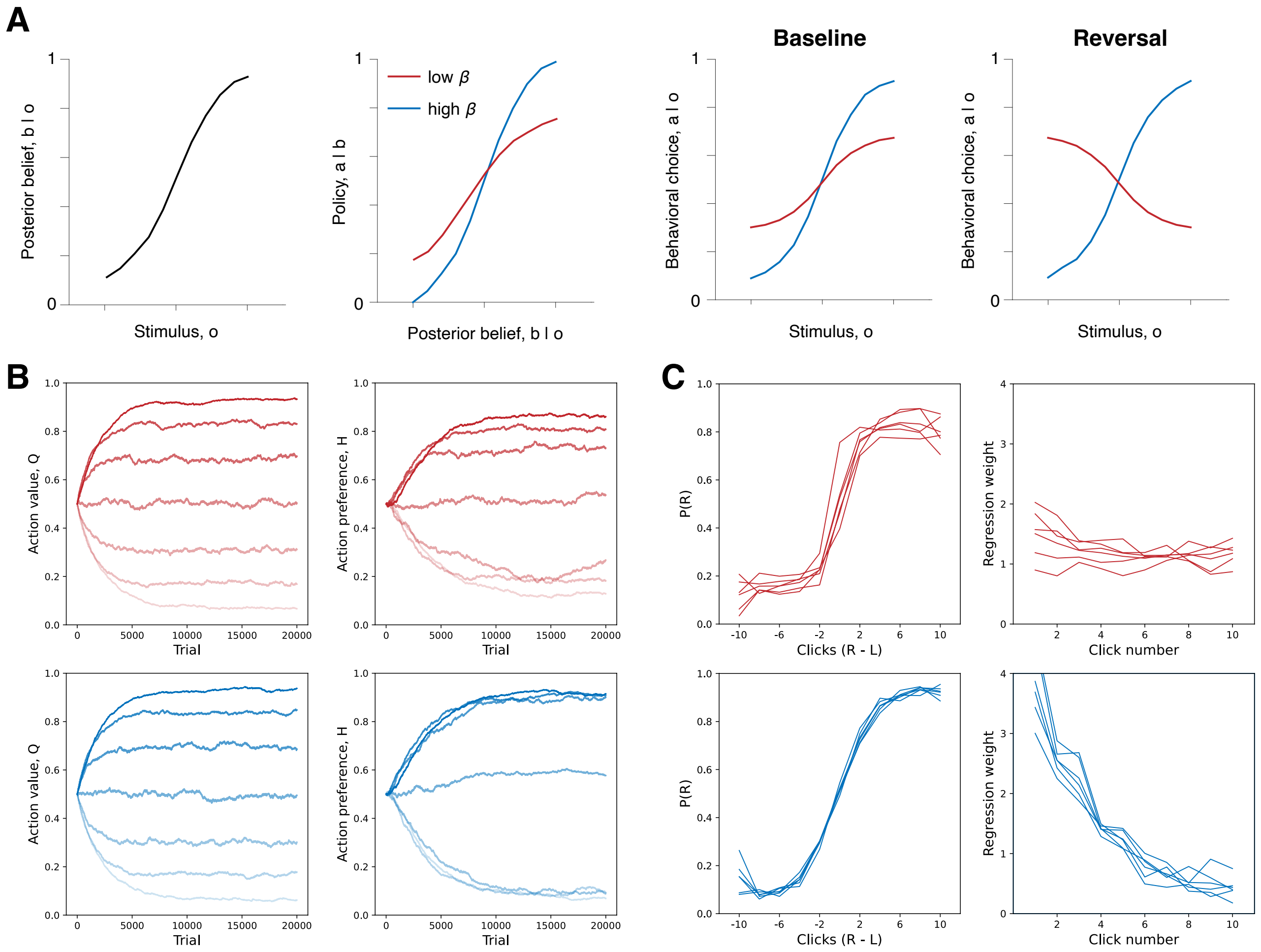


Figure S6: **Habit formation in perceptual learning tasks**. **A.** Left: The posterior belief as a function of the stimulus in an example perceptual inference task. Left middle: The policy i.e., the probability of choosing one of the actions conditioned on the posterior belief under a low (red) and high (blue) degrees of exploitation. Right middle: The probability of choosing one of the actions as a function of the stimulus for the two policies. Right: The probability of choosing one of the actions as a function of the stimulus for the two policies after introducing a reversal in the stimulus-action contingency. **B.** The evolution of action values ($Q$, left) and action preferences ($H$, right) in an evidence accumulation task across learning under conditions of low exploitation (top) or high exploitation (bottom). Different lines correspond to different belief states. Light and dark hues denoting states in which the belief in favor of the proposition that there are more rightward clicks is low and high respectively. **C.** Left: Psychometric curves showing the proportion of rightward response as a function of the difference in rightward and leftward clicks in the stimulus sequence, in models with low (subcritical, top) and high (supracritical, bottom) degree of exploitation. Right: Psychophysical kernels showing the influence of each cue ('click') on the final action in models with low (top) and high (bottom) degree of exploitation.

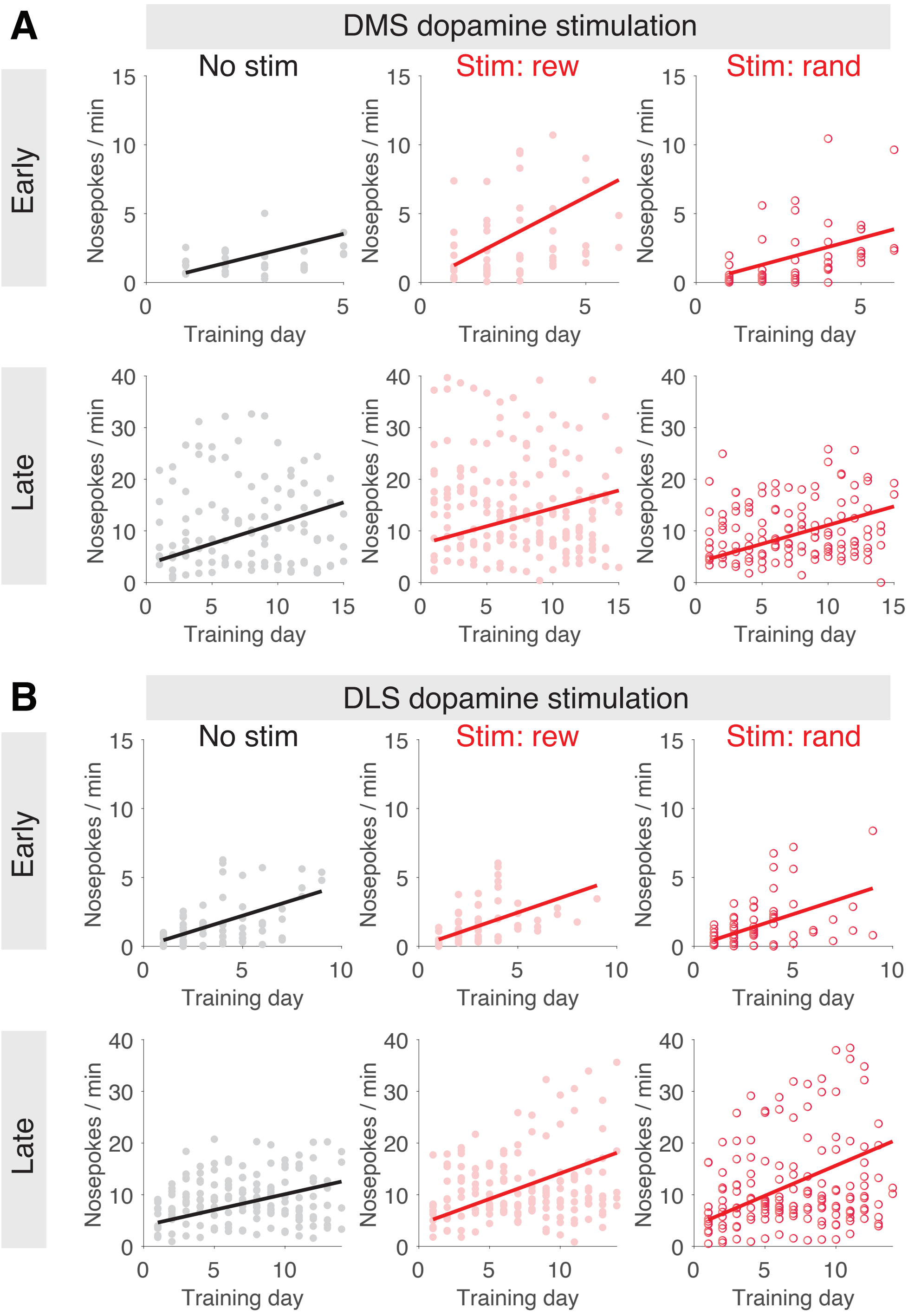


Figure S7: **DMS and DLS dopamine stimulation in mice**. **A.** The evolution of the rate of nose pokes in (Seiler et al., 2022) across early (top row) and late (bottom row) training under conditions in which there was no DMS stimulation (left), stimulation at the time of rewarded nose pokes (middle), and stimulation at the time of random nose pokes (right). Circles denote data from individual mice and solid lines denote the best fit piece-wise linear regression model. **B.** Similar to panels in **A**, but for DLS stimulation.

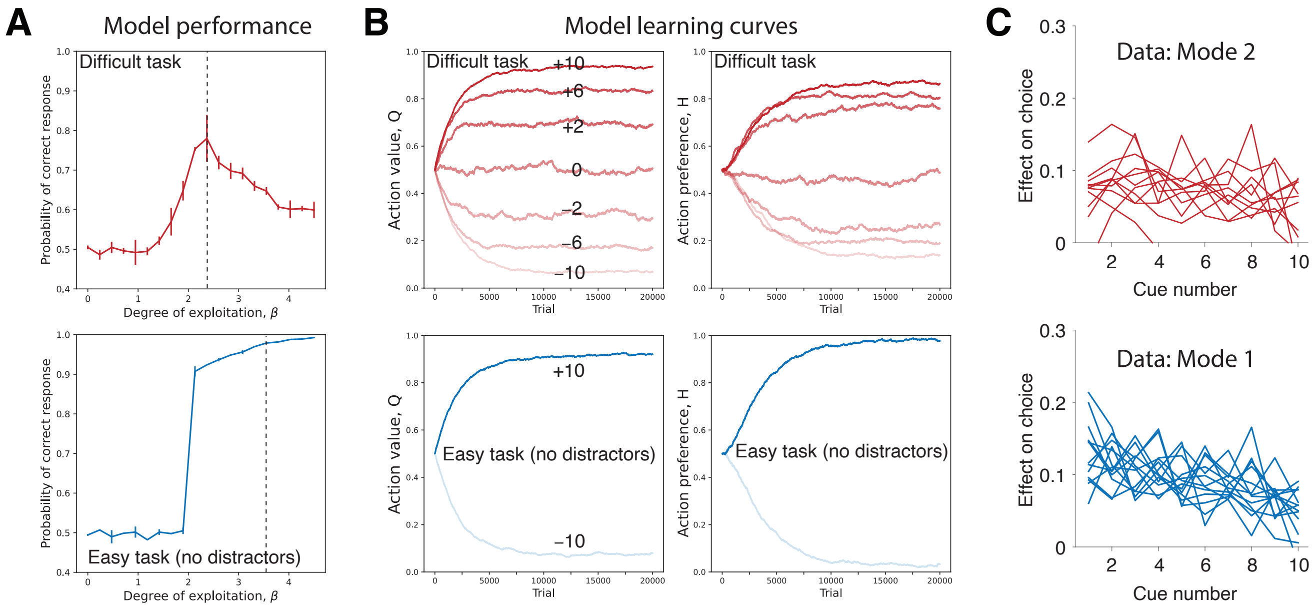


Figure S8: **DMS striatal inhibition in mice**. **A.** Top: Fraction of correct responses of the model as a function of the degree of exploitation ($\beta$), after learning the difficult evidence accumulation task in (Bolkan et al., 2022) i.e., with cues presented on either left/right side during the same trial. Bottom: Similar to the top panel, but for the easy task i.e., no distractor cues. Dashed vertical lines denote the level of exploitation used for simulations in panel **B**. Error bars denote standard error in the mean and 0.5 denotes chance level performance. **B.** The evolution of action values ($Q$, left) and action preferences ($H$, right) across learning in the difficult (top) and easy (bottom) tasks. The numbers overlaid on the lines denote the difference between the number of rightward and leftward cues. **C.** Psychophysical kernels of individual mice during mode 2 (top, $n = 11$) or mode 1 (bottom, $n = 13$). The kernels could not be reliably estimated for a small subset of mice (3 in mode 2; 1 in mode 1) and excluded from these plots.
